# The coordination of spindle-positioning forces during the asymmetric division of the *C. elegans* zygote is revealed by distinct microtubule dynamics at the cortex

**DOI:** 10.1101/638593

**Authors:** H. Bouvrais, L. Chesneau, Y. Le Cunff, D. Fairbrass, N. Soler, S. Pastezeur, T. Pécot, C. Kervrann, J. Pécréaux

## Abstract

In the *Caenorhabditis elegans* zygote, astral microtubules generate forces, pushing against and pulling from the cell periphery. They are essential to position the mitotic spindle. By measuring the dynamics of astral microtubules at the cortex, we revealed the presence of two populations, residing there for 0.4 s and 1.8 s, which correspond to the pulling and pushing events, respectively. Such an experiment offers a unique opportunity to monitor both forces that position the spindle under physiological conditions and study their variations along the anteroposterior axis (space) and the mitotic progression (time). By investigating pulling-force-generating events at the microscopic level, we showed that an anteroposterior asymmetry in dynein on-rate – encoding pulling-force imbalance – is sufficient to cause posterior spindle displacement. The regulation by spindle position – reflecting the number of microtubule contacts in the posterior-most region – reinforces this imbalance only in late-anaphase. Furthermore, we exhibited the first direct proof that the force-generator increasing persistence to pull (processivity) accounts for the temporal control of pulling force throughout mitosis. We thus propose a three-fold control of pulling force, by the polarity, spindle position and mitotic progression. Focusing on pushing force, we discovered a correlation between its density and the stability of the spindle position during metaphase, which strongly suggests that the pushing force contributes to maintaining the spindle at the cell centre. This force remains constant and symmetric along the anteroposterior axis during the division. The pulling one increases in intensity and becomes dominant at anaphase. In conclusion, the two-population study enabled us to decipher the complex regulation of the spindle positioning during cell division.

## INTRODUCTION

During asymmetric division, the position of the mitotic spindle is accurately regulated. Its final position participates in the correct partition of cell fate determinants, which is crucial to ensure faithful division during developmental processes (Gönczy, 2008; Neumüller and Knoblich, 2009; Morin and Bellaïche, 2011; McNally, 2013; Kotak, 2019). Furthermore, its position at the late metaphase controls the pulling-force burst (Bouvrais *et al.*, 2018). In the one-cell embryo of the nematode *Caenorhabditis elegans*, the mitotic spindle is first oriented along the polarity axis and positioned at the cell centre. Then, the spindle is maintained at that position for a few minutes during metaphase. Finally, it is displaced towards the posterior before division (Gönczy, 2008; McNally, 2013). So far, cell-scale investigations revealed the forces at the core of this precise choreography but remained elusive in their regulation. In particular, force generators pull on astral microtubules from the cell cortex and cause the posterior displacement. The force generators are composed of the dynein/dynactin complex, the LIN-5^NuMA^ protein, and the G-protein regulators GPR-1/2^LGN^ and are anchored at the membrane through Gα subunits (Gotta and Ahringer, 2001; Colombo *et al.*, 2003; Srinivasan *et al.*, 2003; Couwenbergs *et al.*, 2007; Nguyen-Ngoc *et al.*, 2007). This trimeric complex generates forces through dynein acting as molecular motor and/or tracking the plus-end of depolymerising microtubule (Schmidt *et al.*, 2005; Kozlowski *et al.*, 2007; Nguyen-Ngoc *et al.*, 2007; O’Rourke *et al.*, 2010; Laan *et al.*, 2012a). Opposite to this cortical pulling, the centring force maintains the spindle at the cell centre during metaphase. Its mechanism is still debated with three major possibilities (Wühr *et al.*, 2009; Wu *et al.*, 2017a): specific regulation of the cortical pulling forces (Tsou *et al.*, 2002; Grill and Hyman, 2005; Kimura and Onami, 2007; Gusnowski and Srayko, 2011a; Laan *et al.*, 2012a); pulling forces generated again by dynein localised at cytoplasmic organelles (Kimura and Onami, 2005; Kimura and Kimura, 2011; Shinar *et al.*, 2011; Barbosa *et al.*, 2017); and cortical pushing forces resulting from the growing of astral microtubules against the cortex (Garzon-Coral *et al.*, 2016; Pécréaux *et al.*, 2016), similarly to the mechanism found in yeast (Tran *et al.*, 2001; Tolic-Nørrelykke *et al.*, 2004). So far, these studies were all based on cell-scale measurements.

How are the cortical pulling and pushing forces regulated and coordinated in space and throughout mitosis? The previous studies approached them separately, resorting to spatial or temporal averages. The cortical pulling forces are asymmetric, because of a higher number of *active* force generators – trimeric complexes engaged in pulling events with astral microtubules – at the posterior-most region of the embryo (Gotta *et al.*, 2003; Grill *et al.*, 2003; Pécréaux *et al.*, 2006a; Nguyen-Ngoc *et al.*, 2007; Rodriguez-Garcia *et al.*, 2018), in response to polarity cues (Grill *et al.*, 2001; Colombo *et al.*, 2003; Tsou *et al.*, 2003; Park and Rose, 2008; Bouvrais *et al.*, 2018). Besides, the physical basis of the progressive increase in the pulling force along the course of the division was inferred from cell-scale measurements, particularly during anaphase, and a molecular mechanism is still missing (Labbé *et al.*, 2004; Pécréaux *et al.*, 2006a; Campbell *et al.*, 2009; Bouvrais *et al.*, 2018). Furthermore, the spatiotemporal regulation of the centring force is still unknown, as well its coordination with opposed pulling force. We here addressed this gap, through analysing the astral microtubules contacting the cortex.

Astral microtubules are involved in generating all these forces. These semi-flexible filaments emanate from the spindle poles. They are dynamic, switching alternatively from growing to shrinking and back, at the catastrophe and rescue rates, respectively (Mitchison and Kirschner, 1984). At the cortex, astral microtubules can be in three different states: shrinking in coordination with cortex-anchored dynein that generates pulling force (Gonczy *et al.*, 1999; Dujardin and Vallee, 2002; Grishchuk *et al.*, 2005; Gusnowski and Srayko, 2011b; Laan *et al.*, 2012a; Rodriguez-Garcia *et al.*, 2018); pushing by growing against the cortex, likely helped by stabilising associated proteins like CLASP (Faivre-Moskalenko and Dogterom, 2002; Dogterom *et al.*, 2005; Howard, 2006; Espiritu *et al.*, 2012); or stalled, clamped possibly by dynein tethering or other proteins (Labbé *et al.*, 2003; Sugioka *et al.*, 2018). Do the microtubule dynamics, especially their cortical residence times, reflect these different states? Interestingly, dynein tethering delays microtubule catastrophe, as shown *in vitro* and by computational studies (Hendricks *et al.*, 2012; Laan *et al.*, 2012a). Oppositely, the larger the pushing force, the smaller the residence time (Janson *et al.*, 2003). In *C. elegans* embryo, microtubules involved in pulling or pushing forces may display different cortical residence times (Pécréaux *et al.*, 2006a; Pécréaux *et al.*, 2016). They could thus reveal the corresponding force-generating events. For instance, previous studies uncovered anteroposterior variations in residence time. The microtubules would be more dynamic (lower lifetime) at the posterior cortex compared to the anterior (Labbé *et al.*, 2003; Sugioka *et al.*, 2018). However, the reported residence times are strikingly different between studies (Labbé *et al.*, 2003; Kozlowski *et al.*, 2007; O’Rourke *et al.*, 2010; Hyenne *et al.*, 2012; Schmidt *et al.*, 2017; Sugioka *et al.*, 2018). How the microtubule residence times evolve throughout mitosis is, however, yet to be studied. Indeed, the short duration of these cortical fluorescent spots of labelled microtubules (a few frames) and the low signal-to-noise ratio of the images made resolving both time and space variations hard until now. Recent developments in microscopy and image-processing tools call for revisiting this problem (Chenouard *et al.*, 2014; Kervrann *et al.*, 2015).

Beyond imaging improvements, the statistical analysis of the durations of microtubule tracks at the cortex – resulting from the detection of the same fluorescent spots over several images – could also be significantly refined in contrast to the classic fit with a mono-exponential distribution (Kozlowski *et al.*, 2007; Sugioka *et al.*, 2018). In particular, we here aim to distinguish several co-existing dynamical behaviours. Thus, we fitted the experimental distribution of the track durations with finite-mixture-of-exponential models and then used an objective criterion to choose the best one. Such an approach, although delicate, benefits from developments in applied mathematics (Grinvald and Steinberg, 1974; James and Ware, 1985; Vieland and Hodge, 1998; Jae Myung *et al.*, 2000; Turton *et al.*, 2003). Furthermore, in our case, the microtubule residence times could last only a few tenths of a second, i.e. a few frames. The discrete nature of the residence-time histogram calls for specific analysis as performed in photon counting experiments. This field has designed appropriate fitting strategies that offer a firm starting point to analyse microtubule dynamics (Maus *et al.*, 2001; Turton *et al.*, 2003; Nishimura and Tamura, 2005; Laurence and Chromy, 2010).

In the present paper, we aim to study the spatiotemporal regulation of the spindle-positioning forces during the first mitotic division of the *C. elegans* embryo. To do so, we measured the microtubule dynamics at the cortex. We designed the *DiLiPop assay* (Distinct Lifetime subPopulation) to disentangle several microtubule populations distinct by their cortical residence times. We found two of them, which we could associate with different microtubule functions. Equipped with this assay, we could investigate in time and space, and at the microscopic level, the regulation of the forces positioning the spindle during mitosis. We directly measured the force-generator processivity increase that accounts for the pulling force regulation throughout mitosis. We showed that the three controls of pulling force (by polarity, spindle position and mitotic progression) act independently. We also identified which mechanism maintains the spindle at the cell centre during metaphase. Finally, we suggest how the two cortical forces, pushing and pulling, coordinate in space and time.

## RESULTS

### The Distinct Lifetime (sub)Population assay reveals two populations of microtubules at the cortex

To investigate the regulation of the forces exerted on the mitotic spindle during the first mitosis of the *C. elegans* zygote, we set to measure the dynamics of astral microtubules at the cortex. The microtubules were entirely fluorescently labelled using YFP::α-tubulin to view them in all their states. We performed spinning disk microscopy at the cortical plane, at 10 frames per second similarly to (Bouvrais *et al.*, 2018) (Supp. Text §1.1.1). When the microtubules contacted the cortex end-on, they appeared as spots (Movie S1). The high frame rate needed to resolve the brief cortical contacts led to images with a low signal-to-noise ratio (Figure 1A, top). We mitigated this issue by denoising using the Kalman filter (Figure 1A, middle) (Kalman, 1960). We then tracked the microtubule contacts using the u-track algorithm (Figure 1A, bottom, Table S1, Supp. Text §1.1.2) (Jaqaman *et al.*, 2008). This image-processing pipeline is further named *KUT*. We estimated that it enabled us to capture at least 2/3 of the microtubule contacts (Figure S1A) by comparing with electron tomography (Redemann *et al.*, 2017).

**Figure 1:**
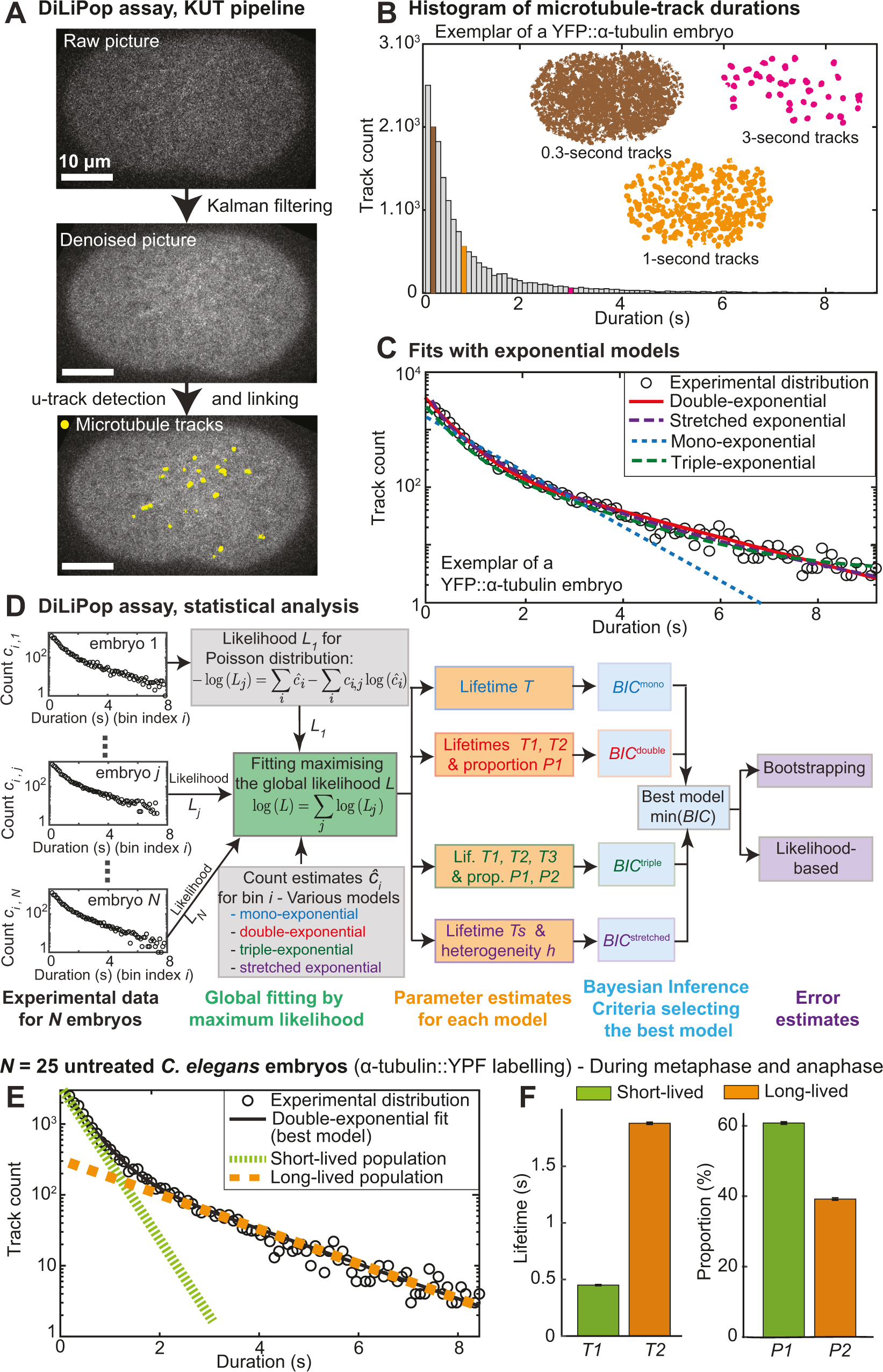
Microtubule dynamics at the cortex of the *Caenorhabditis e l egans* embryo encompass two distinct residence-time behaviours during the first zygotic division. (**A-D**) The typical workflow of the DiLiPop (<u>Di</u>stinct <u>Li</u>fetime sub<u>Pop</u>ulation) assay disentangles microtubule populations. (**A**) Exemplar KUT analysis of a one-cell embryo. Bright spots are the plus-ends of YFP::α-tubulin labelled microtubules (top) (Movie S1). They are enhanced after denoising by a Kalman filter (middle). The trajectories of the microtubules (yellow lines) are obtained using the u-track algorithm (bottom) (Supp. Text, §1.1.2). The parameters used are in Table S1. (**B**) Experimental distribution of the microtubule-track durations for a typical untreated embryo imaged from nuclear envelope breakdown (NEBD) to late anaphase at 10 frames per second. (Insets) Spatial distributions of the tracks lasting 0.3 s (brown), 1 s (orange), and 3 s (pink). (**C**) The above experimental distribution (open circles) was fitted using various exponential models: (dashed blue line) mono-exponential, (plain red line) double-exponential, (dashed green line) triple-exponential, and (dashed purple line) stretched exponential (Supp. Text, §1.2.1). (**D**) Flow diagram of the advanced statistical analysis used in the DiLiPop assay (Supp. Text, §1.2). (White boxes) Exemplar distributions (histograms), depicting the count *c*_*i,j*_ per duration-bin (indexed by *i*), *j* indexing the embryo. (Grey shadings) The experimental distributions of the microtubule-track durations for *N* embryos were individually fitted using different exponential models (Supp. Text, §1.2.1) and assuming a Poisson distribution (Supp. Text, §1.2.2). (Green shading) We maximised the global likelihood *L*, computed as the product of embryo likelihoods *L*_*j*_ (Supp. Text, §1.2.3). (Orange shadings) We thus obtained the model parameters for each studied model. (Blue shadings) The best model was selected as the one minimising the Bayesian Inference Criterion (*BIC*) (Supp. Text, §1.2.4). (Purple shadings) We estimated the standard deviations on the best-model parameters by using either a bootstrap approach (Figure S1D) or the likelihood-based confidence intervals (Figure S1E) (Supp. Text, §1.2.5). (**E**) Microtubule-track durations of *N* = 25 untreated embryos (same condition as in B-C) were subjected to DiLiPop global fit. The best-fitting model was the double exponential (black line). Dotted green and dashed orange lines highlight the separate contributions of each exponential component, respectively short- and long-lived. The *BIC* values for each model are reproduced in Table S2. **(F)** Corresponding fit parameters and error bars by bootstrapping.

We computed the duration distributions of the microtubule tracks for each embryo separately (to avoid averaging artefacts) (Figure 1B). When all the microtubules have the same catastrophe rate, this distribution follows an exponential decay (Kozlowski *et al.*, 2007; Floyd *et al.*, 2010). However, we also envisaged that multiple mechanisms involving microtubules are superimposed, leading to distinct catastrophe rates. Therefore, we fitted the distribution with finite-mixture-of-exponential models, in particular double- and triple-exponential models (Supp. Text §1.2.1). The double-exponential appeared to fit the duration-distribution better, suggesting that we observed at least two populations of microtubules contacting the cortex of *C. elegans* embryo, distinct by their residence times (Figure 1C). These populations may offer the opportunity to visualise the various force-generation mechanisms. To finely characterise them, we implemented an advanced statistical analysis of the track-duration distribution (Figure 1D), described in details in Supp. Text §1.2. In a nutshell, because we fitted a histogram with few counts in some bins, we modelled the data point errors using a Poisson law. We designed the objective function correspondingly to fit the histogram (Figure 1D, grey shading) (Supp. Text §1.2.2) (Laurence and Chromy, 2010). To distinguish between multiple populations within each embryo from a single population per cell with parameters varying between embryos, we fitted each embryo individually. However, to gain certainty, we imposed the same model parameters on each embryo of a dataset, by global fitting, i.e. maximising the product of the embryo-wise likelihoods (Figure 1D, green) (Supp. Text §1.2.3) (Beechem, 1992). We performed an unbiased selection of the best mixture-of-exponential model using the Bayesian Inference Criterion (*BIC*) (Figure 1D, blue) (Supp. Text §1.2.4) (Schwarz, 1978). Finally, we computed the confidence intervals on the fitted parameters using bootstrapping (Figure 1D, purple; Figure S1D) (Supp. Text §1.2.5) (Efron and Tibshirani, 1993). We validated this approach using the likelihood ratio (Figure S1E) (Bolker, 2008; Agresti, 2013). Applying this approach to untreated embryos of *C. elegans*, we found two populations within the microtubules residing at the cortex (Figure 1EF, Table S2). Their distinct dynamics suggest that the microtubules could be involved in two different mechanisms.

We firstly ensured that our complex pipeline could not create the two dynamically distinct populations through artefacts. We built images containing particles with a single dynamical behaviour (Figure S2A, black; Table S3; Material and Methods) (Costantino *et al.*, 2005). By DiLiPop analysis, we recovered a single population with the correct lifetime (Figure S2A, red). In contrast, a similar simulation with two dynamical populations led to the double-exponential as best model and accurate parameters (Figure S2B, blue). Overall, the KUT image-processing pipeline does not cause artefacts. However, and to gain certainty, we repeated the analysis of *in vivo* data using an image-processing pipeline based on different hypotheses. This pipeline, named *NAM*, encompasses the ND-SAFIR denoising (Figure S2F, middle) (Boulanger *et al.*, 2010), the ATLAS spot-detecting (Basset *et al.*, 2015), and the MHT linking (Multiple Hypothesis Tracker) (Figure S2F, right) (Chenouard *et al.*, 2013), with settings listed in Table S4 (Supp. Text §1.1.3). Applied to untreated *C. elegans* embryos, the NAM pipeline combined to DiLiPop statistical analysis recovered the two populations distinct by their dynamics. Furthermore, the lifetimes are close to the ones obtained using the KUT pipeline (Figure S2C). We therefore excluded that the two dynamically distinct populations could be artefactual.

Before investigating the biological relevance of these populations, we wondered whether there might be even more than two. We reasoned that the number of data points, typically ∼20 000 microtubule tracks per embryo, may be insufficient to support a triple-exponential model. We addressed this question *in silico* and simulated distributions of microtubule-track durations creating “*simulated embryos*”, with three dynamical populations of lifetimes 0.4 s, 1.5 s and 4 s, and proportions set to 55%, 40% and 5%, respectively. These values correspond to experimental estimates on untreated embryos (Table S2). To mimic the experimental conditions of untreated embryos, we generated “*fabricated datasets*” composed of 25 simulated embryos and analysed them using the DiLiPop assay. We repeated 10 times this simulation procedure to get certainty about the results. We further considered only the sample sizes, for which a majority of fabricated datasets led to the simulated model, here triple exponential, being the best model according to Bayesian criterion. Among valid conditions, we averaged the recovered model parameters over the datasets, where the recovered best model was correct. It suggests that 20 000 tracks per embryo were necessary and also sufficient to support the triple-exponential model if applicable (Figure S3A). We reckoned that a third and very-long-lived population might be in such a low proportion that the amount of experimental data did not allow identifying it. Keeping with our *in silico* approach and using 20 000 tracks per embryo, we fixed the short-lived proportion to 55% and very-long-lived one from 2.5% to 10%. We found that 5% is enough to support the triple-exponential model (Figure S3B). We concluded that in untreated embryos, there is a less than 5% of very-long-lived population of astral microtubules.

We wondered whether two well-defined microtubule populations exist or whether the numerous molecular motors and MAPs could lead to a broadly-varying microtubule residence times. We modelled this latter case using a stretched exponential (Lee *et al.*, 2001; Siegel *et al.*, 2001). Such a model was not the best using the untreated embryo data (Table S2). However, we again asked whether the amount of experimental data was sufficient, using *in silico* approach. We simulated microtubule durations displaying a stretched exponential behaviour of lifetime 0.1 s and heterogeneity parameter 2.2, which are the experimental estimates on untreated embryos (Table S2). We found that 500 tracks per embryo were sufficient (Figure S3C). Because we had far more tracks in experimental data, we concluded that the two dynamical behaviours measured *in vivo* truly correspond to two populations of microtubules.

Being confident in the biological origin of the two microtubule-populations, we next asked whether it reflects truly the force-generating events. Alternatively, the labelling or variations in the dosage of tubulin paralogs within the microtubules could account for our observations (Wright and Hunter, 2003; Honda *et al.*, 2017). As an alternative to the labelling used above, YFP::TBA-2^α-tubulin^, we repeated our experiment using GFP::TBB-2 ^β-tubulin^ (Figure S2D1) and measured two populations of microtubules at the cortex, with similar lifetimes (Figure S2D2). The change in lifetimes was larger for the long-lived population and could originate from distinct dye-brightness or sensitivity to bleaching. These differences are comparable to the one observed when changing the image processing pipeline (Supp. Text §1.1.3) We concluded that the two microtubule populations were likely to reflect distinct force-generating events.

Finally, we wondered whether such two microtubule-populations exist beyond C. *elegans*. We investigated the microtubule dynamics at the cortex in a cousin nematode species, *Caenorhabditis briggsae*, where β-tubulin was labelled. We again observed two populations distinct by their dynamics (Figure S2E). We concluded that these two populations are not a peculiarity of *C. elegans* embryo. Overall, by viewing the microtubule contacts in all their states, we measured two populations at the cortex. Because they are dynamically distinct, a possible interpretation was that they reflected pulling and pushing force-generating events. Indeed, pushing microtubules are likely to reside longer to contribute to centring (Garzon-Coral *et al.*, 2016; Pécréaux *et al.*, 2016; Howard and Garzon-Coral, 2017), while short residence times could correspond to events of pulling by dynein, proposed to last about 0.5 s (Pécréaux *et al.*, 2006a; Rodriguez-Garcia *et al.*, 2018).

### The short- and long-lived microtubules correspond to events of pulling from and pushing against the cortex

More dynein engaged in pulling on the posterior side causes the cortical pulling-force imbalance and the spindle posterior-displacement during the anaphase of the asymmetric division of the nematode zygote (Grill *et al.*, 2003; Redemann *et al.*, 2010; Rodriguez-Garcia *et al.*, 2018). However, we reckoned that the distribution of short-lived and long-lived contacts might be different. We refined the DiLiPop assay to map the cortical contacts along the anteroposterior axis (AP axis) within each population. By selecting biologically relevant regions and time-blocks as small as possible, we guaranteed enough data to detect two populations accurately (Supp. Text §1.3, Figure S4). We applied this analysis to untreated embryos and recovered the high contact-density ridgelines, for both populations, as previously reported (Bouvrais *et al.*, 2018) (Figure 2A). About 50 s before the anaphase onset, the instantaneous contact-density of the short-lived population increased posteriorly and became asymmetric (Figure 2A1). The short-lived microtubules are particularly enriched in the region where the force generators are active and which extends from 70% to 100% along the AP axis (Krueger *et al.*, 2010; Bouvrais *et al.*, 2018). In contrast, the long-lived contact density remains symmetric, with a slight posterior enrichment in late anaphase, expected because the spindle displaces towards the posterior (Figure 2A2) (Bouvrais *et al.*, 2018). The specific polarisation of the short-lived population suggests that the corresponding microtubule contacts reveal pulling force-generating events.

**Figure 2:**
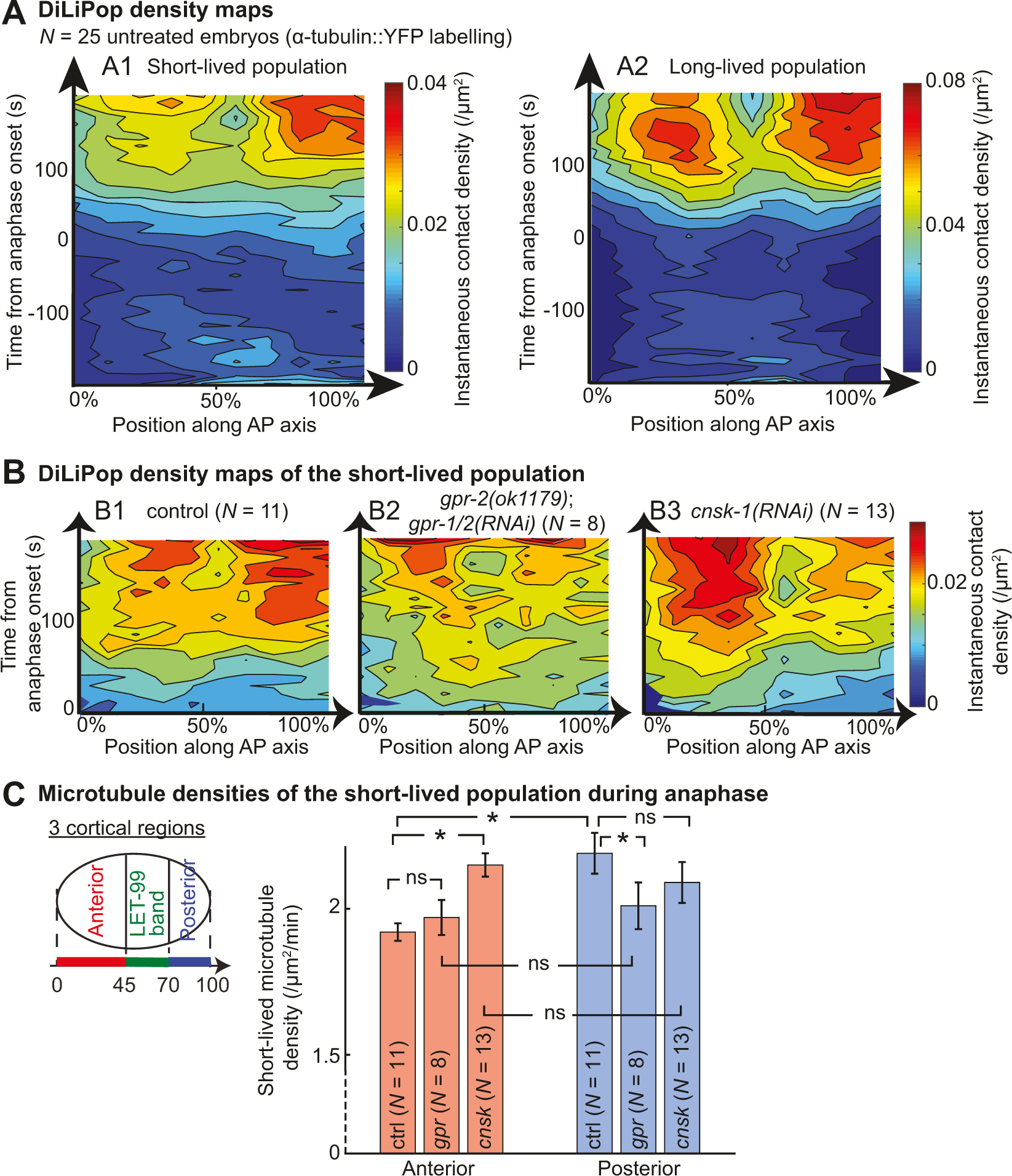
Microtubules pulling from the cortex belong to the short-lived population. (**A**) DiLiPop density maps, computed for a dataset of *N* = 25 untreated α-tubulin-labelled embryos, show the instantaneous distributions of (A1) the short-lived and (A2) the long-lived contacts along the anteroposterior axis (AP axis), during metaphase and anaphase. The DiLiPop mapping attributes each contact to a population (Supp. Text, §1.3). We used 3 regions and 60-s time-blocks since 25 simulated embryos featuring 350 tracks each are enough to ensure detecting two populations (Figure S4B). The heat map is computed by averaging the mapped contacts within 10 regions of equal width along the AP axis and over a 10-s running window for each embryo. These maps were then averaged over the dataset (Supp. Text, §1.3). (**B**) Short-lived-contact density maps of (B1) *N* = 11 control embryos, (B2) *N* = 8 *gpr-2(ok1179);gpr-1/2(RNAi)-*treated embryos and (B3) *N* = 13 *cnsk-1(RNAi)-*treated embryos. To determine the characteristics of the two populations, we used 3 regions and the whole anaphase, based on estimating track count requirement by analysing 8 simulated embryos featuring 1000 tracks each as above (Figure S5A). (**C**) Corresponding comparisons of the short-lived microtubule densities in the anterior region (0 - 45% of AP axis, red) and posterior-most region (70 - 100% of AP axis, blue). Error bars are the standard deviations (SD) obtained by bootstrapping (Supp. Text, §1.2.5). Star indicates significant differences (Student’s *t*-test). For concision, *gpr* stands for *gpr-2(ok1179);gpr-1/2(RNAi)* and *cnsk* for *cnsk-1(RNAi)*.

To further support this result, we genetically decreased or increased cortical pulling forces and observed the spatial distribution of the short-lived microtubule density (Rodriguez-Garcia *et al.*, 2018). Firstly, we depleted GPR-1/2^LGN^, the well-established force generator regulator (Colombo *et al.*, 2003; Grill *et al.*, 2003; Pécréaux *et al.*, 2006a; Nguyen-Ngoc *et al.*, 2007). We used *gpr-2(ok1179)* mutant embryos with *gpr-1/2(RNAi)* treatment to ensure a strong depletion and computed the DiLiPop-map. We observed a significant reduction in the short-lived microtubule density in the posterior-most region compared to the control during anaphase (Figure 2B1-2, 2C). It led to cancelling out the asymmetric distribution of this population. In similar conditions, we imaged at the spindle plane and tracked the spindle poles in *N* = 8 embryos. We observed a loss of spindle-pole oscillation and a strong reduction of the spindle posterior-displacement as reported previously (Colombo *et al.*, 2003; Pécréaux *et al.*, 2006a) (Figure S6D1-2). Since the short-lived-population distribution strongly depends on GPR-1/2, the corresponding microtubules are likely to contribute to generating pulling force.

Secondly, we performed the converse experiment, enriching the force generators anteriorly through a *cnsk-1(RNAi)* treatment (Panbianco *et al.*, 2008). We observed a significant increase in the short-lived microtubule density anteriorly, compared to controls (Figure 2B3, 2C), consistent with previous observation using labelled dynein (Rodriguez-Garcia *et al.*, 2018). We also observed a slight decrease in the short-lived densities at the posterior-most region attributed to the anterior displacement of the spindle (Figure S6D1,D3) (Bouvrais *et al.*, 2018; Rodriguez-Garcia *et al.*, 2018). Under the same conditions and at the spindle plane, we observed a clear increase of centrosome oscillation amplitudes anteriorly 4.0 ± 0.2 µm (*N* = 11 embryos) compared to 2.2 ± 0.1 µm in control embryos (two-tailed Student’s *t*-test: *p* = 9×10^−3^, *N =* 7 control embryos). We also measured increased oscillations at the posterior pole, although not significantly, with a peak-to-peak amplitude of 5.4 ± 0.1 µm compared to 5.1 ± 0.2 µm (two-tailed Student’s *t*-test: *p* = 0.62). It confirms the significant increase in pulling forces mostly at anterior (Panbianco *et al.*, 2008). Overall, the short-lived microtubule density correlates with both the cortical force intensity and the number of active force generators, supporting our interpretation of the short-lived population.

We reckoned that the long-lived population might correspond to microtubules pushing against the cortex. To challenge this idea, we impaired microtubule growth by depleting the promoting factor ZYG-9^XMAP215^ by RNAi. We then performed a DiLiPop analysis during metaphase without splitting into regions to gain accuracy and since the long-lived population is not polarised We found two populations, but interestingly, only the long-lived microtubules had their lifetime significantly reduced while the short-lived one was unaltered (Figure 3A1-2, green). It was consistent with the reported activity of ZYG-9 (Bellanger and Gönczy, 2003; Srayko *et al.*, 2003; Brouhard *et al.*, 2008). It supported our hypothesis that the long-lived population accounts for pushing microtubules. Under the same conditions and at the spindle plane, we observed a reduction of the metaphase spindle length before elongation, which reads 8.7 ± 0.7 µm upon *zyg-9(RNAi)* (*N* = 8) compared to 10.2 ± 0.9 µm in control embryos (*p* = 3.7×10^−3^, *N* = 7), as expected (Srayko *et al.*, 2003). To strengthen the link between the long-lived population and the growing microtubules, we partially depleted the microtubule-depolymerising kinesin KLP-7^MCAK^ by a hypomorphic RNAi treatment. The DiLiPop analysis revealed no significant change in the lifetime of the long-lived population during metaphase (Figure 3A1, blue). We also found that the short-lived population displayed a slightly increased residence time (Figure 3A2, blue) consistent with the increased pulling forces previously reported (Grill *et al.*, 2001; Gigant *et al.*, 2017). When imaging at the spindle plane during anaphase, we measured a faster spindle elongation equal to 0.156 ± 0.020 µm/s upon *klp-7(RNAi)* (*N* = 9) compared to 0.102 ± 0.026 µm/s for the control embryos (*p* = 1.8×10^−5^, *N* = 13), as expected (Grill *et al.*, 2001; Gigant *et al.*, 2017). We concluded that the long-lived population reflects specifically microtubule growing against the cortex, leading to pushing force. Importantly, both KLP-7 and ZYG-9 depletion experiments are consistent with associating short-lived population with pulling force generation.

**Figure 3:**
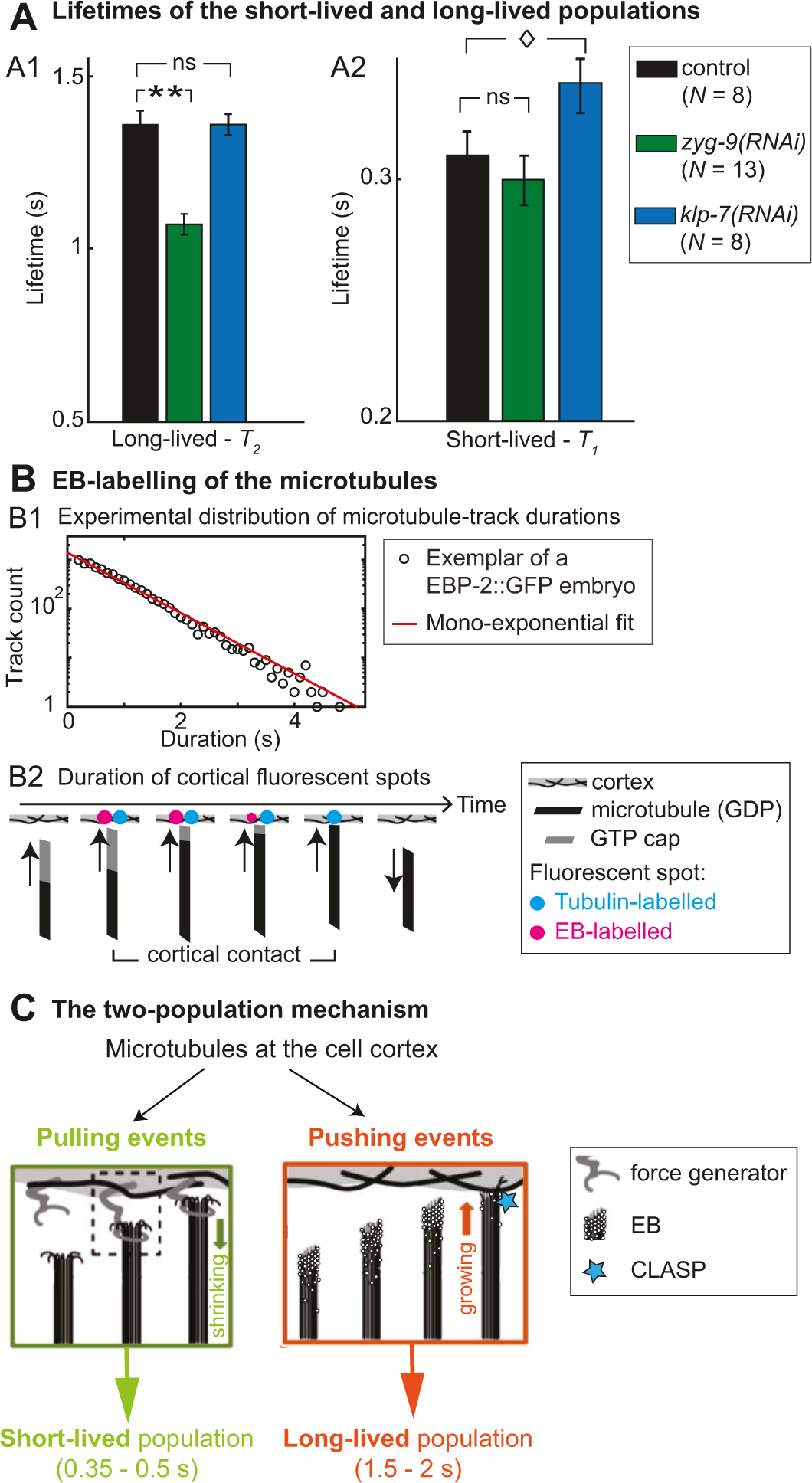
The microtubules pushing against the cortex belong to the long-lived population. **(A)** DiLiPop analysis of the microtubule dynamics at the cortex during metaphase, in *N* = 13 *zyg-9(RNAi)-*treated embryos (green), *N* = 8 *klp-7(RNAi)-* treated embryos (blue), and *N* = 8 control embryos (black). We compared the lifetimes of (A1) the long-lived and (A2) the short-lived populations. Error bars are the standard deviations (SD) obtained by bootstrapping (Supp. Text, §1.2.5). Stars or diamond indicate significant differences (Student’s *t*-test). (**B**) (B1) Experimental distribution of the microtubule-track durations for a typical EB-labelled embryo. Over *N* = 9 embryos, the distributions were best fitted by a mono-exponential, with a lifetime equal to 0.64 s. (B2) Schematic highlighting the putative mechanism causing different cortical residence times upon EB- (pink) and tubulin-labellings (light blue). (**C**) The two-population mechanism: the short-lived microtubules account for pulling-force generating events, while the long-lived ones for pushing-force generating events.

To better distinguish the two populations, we set to label specifically the growing microtubules using an EBP-2::GFP strain. We found a single microtubule population with a lifetime intermediate between the two obtained using YFP::α-tubulin labelling (Figure 3B1). On the one hand, we could attribute this latter result to a direct effect of EBP-2 over-expression, which would alter microtubule dynamics, as seen in other organisms (Duellberg *et al.*, 2016). On the other hand, the microtubule can reside at the cortex and push against it with a reduced GTP cap resulting in a loss of EBP-2::GFP signal but not YFP::α-tubulin one (Figure 3B2). Indeed, *in vitro* experiments found a delay between the CAP disappearing and catastrophe (Bieling *et al.*, 2007; Kozlowski *et al.*, 2007; Zanic *et al.*, 2009). Indeed, proteins like CLASP or even dynein can stabilise the microtubule (Espiritu *et al.*, 2012; Laan *et al.*, 2012b). In a broader take, we suggest that the long-lived population reflects specifically microtubules pushing against the cortex (Figure 3C, orange). Meanwhile, perturbations of cortical-pulling-force level or distribution are visible on the short-lived microtubules, relating these latter to the pulling force-generating events (Figure 3C, green).

### The asymmetric dynein on-rate sets the final spindle position independently from positional control and mitotic progression

Multiple mechanisms regulating the cortical pulling forces were proposed. Monitoring them through DiLiPop offered an unparalleled opportunity to investigate the links between these controls, termed polarity, positional and temporal (mitotic progression). Indeed, others and we suggested that mitotic progression is the first regulation through force-generator off-rate, the inverse of the processivity, i.e. the persistence of the force generators to pull on microtubule before detaching (Labbé *et al.*, 2004; Pécréaux *et al.*, 2006a; McCarthy Campbell *et al.*, 2009; Bouvrais *et al.*, 2018). We also proposed that a higher dynein-microtubule on-rate at the posterior cortex compared to the anterior one causes the cortical pulling-force imbalance. It reflects the polarity and accounts for the spindle posterior displacement (Fielmich *et al.*, 2018; Rodriguez-Garcia *et al.*, 2018). This on-rate could be the binding rate of force generator dynein to the microtubules or its engaging rate, i.e. the initiation of a motor run to exert a pulling force. Concurrently, we also reported the regulation of these same forces by the position of the spindle itself (Bouvrais *et al.*, 2018).

We firstly investigated the link between polarity and temporal control. To do so, we compared the anterior (0-45% of AP axis) and posterior-most (70-100% of AP axis) regions over time, using in particular the Wilcoxon signed rank test. We measured the short-lived population since it corresponds to the pulling force. We observed that the asymmetry in the short-lived microtubule density built up along mitosis (Figure 4A1,A3,B1) in contrast to the lifetime that remained mostly symmetric (Figure 4A2,A4,B2). It shows that pulling force imbalance exists from at least early metaphase. It is consistent with dyneins being denser on posterior but persisting same times on both sides (Rodriguez-Garcia *et al.*, 2018). It may suggest that force polarisation is independent of the mitotic progression.

**Figure 4:**
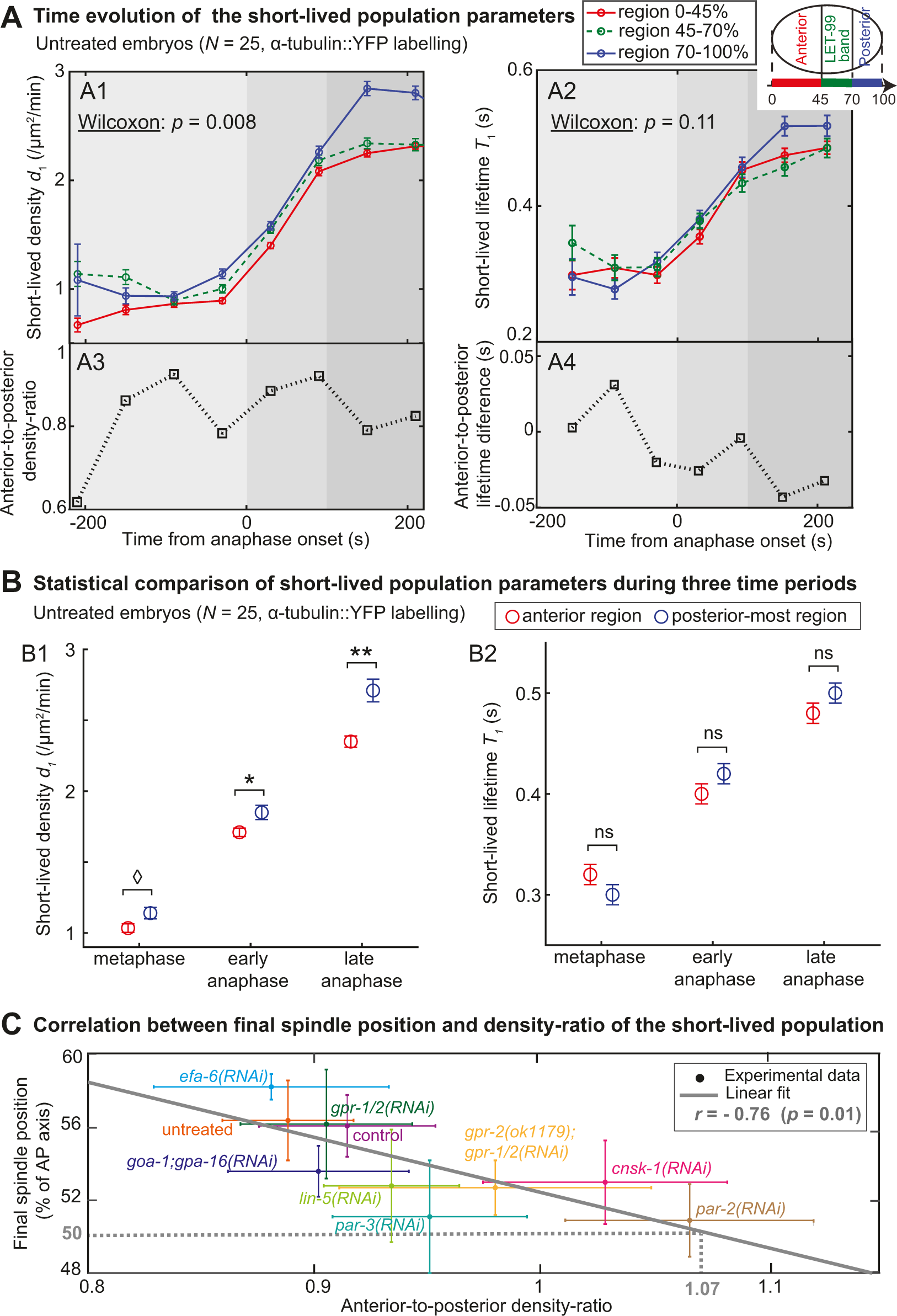
An asymmetry in the short-lived microtubule-density ratio is sufficient to cause the posterior displacement of the spindle. (**A**) Evolution of the short-lived population parameters during metaphase and anaphase: (A1) microtubule densities and (A2) lifetimes, in (red) the anterior region, (green) the lateral LET-99 band and (blue) the posterior-most region. These regions are depicted in the schematics at the top right. We analysed the same embryos as in Figures 2A, 6A and S10AB, i.e. *N* = 25 untreated α-tubulin-labelled embryos, using 60-s time-blocks. Below each plot, either (A3) the anterior-to-posterior density ratio or (A4) the anterior-to-posterior lifetime difference is plotted. Standard deviations were computed by bootstrapping (Supp. Text, §1.2.5). We found a significant difference between the time-series of anterior and posterior-most regions for the densities, as supported by the Wilcoxon signed-rank test, but not for the lifetimes. The grey shadings depict, from lighter to darker, the three time-periods, namely metaphase (the 210 s before anaphase onset), early anaphase (the 100 s after anaphase onset), and late anaphase (from 100 s to 210 s after anaphase onset). (**B**) We analysed the same quantities comparing the two extreme regions and reducing the time resolution to the above three time-periods for the sake of accuracy. Stars or diamond indicate significant differences (Student’s *t*-test). (**C**) Final spindle position obtained by imaging the same strain at the spindle plane, plotted against the anterior-to-posterior density ratio for the short-lived population, assessed during the whole anaphase. The grey line depicts the Pearson anticorrelation. The density-ratio was varied by depleting various proteins: *par-3(RNAi)* (*N* = 10 embryos acquired at the cortex and *N* = 13 at the spindle plane, further written 10/13), *par-2(RNAi)* (*N* = 9/16), *gpr-2(ok1179);gpr-1/2(RNAi)* (*N* = 8/8), *cnsk-1(RNAi)* (*N* = 13/9), *lin-5(RNAi)* (*N* = 13/14), *goa-1;gpa16(RNAi)* (*N* = 12/9), *gpr-1/2(RNAi)* (*N* = 11/6), *efa-6(RNAi)* (*N* = 10/11), control embryos *N* = 11/10, and untreated embryos *N* = 25/9. Error bars are the standard deviations. The dotted grey line indicates the short-lived density-ratio estimated from the linear regression for a centred final position of the spindle.

To further explore the link between these two controls, we treated either wild-type embryos by RNAi against *lin-5*, or *gpr-2(ok1179)* mutant embryos by RNAi against *gpr-1/2*, to symmetrise the pulling dynein density. We observed that the short-lived density of microtubules became symmetric upon both treatments (Figures S6A2,B; S7A2,B). Interestingly, the lifetimes of the short-lived population were not affected, indicating that the reduction of force imbalance was likely independent of the control of processivity, i.e. mitotic progression (Figure S8A,B2). To strengthen our hypothesis, we treated embryos using *goa-1;gpa-16(RNAi)*. This protein is also involved in cortical pulling force and may anchor the trimeric complex (Gotta and Ahringer, 2001; Afshar *et al.*, 2004; Afshar *et al.*, 2005; Park and Rose, 2008). We observed, as expected, a reduced asymmetry of the short-lived densities (Figures S7A3,B). The short-lived microtubule lifetimes upon *goa-1;gpa-16(RNAi)* were similar to control ones (Figure S8B3). In the same conditions, at the spindle plane, we observed a reduced spindle posterior displacement and a suppression of oscillations in these 3 conditions (Figure S6D1-2, S7C). On the one hand, *lin-5(RNAi)* treatment was hypomorphic, because the corresponding protein is involved in earlier processes (van der Voet *et al.*, 2009); on the other hand, *RNAi* targeting two genes, *goa-1* and *gpa-16* is knowingly less efficient. In conclusion, it indicates that the mitotic progression control through force generator processivity acts independently from polarity control through GPR-1/2 posterior enrichment and dynein on-rate.

We reckoned that the regulation of microtubule cortical residence times could be separated from the trimeric complex, but still under the control of polarity proteins PAR-2 and PAR-3 (Labbé *et al.*, 2003; Sugioka *et al.*, 2018). As expected, *par-3(RNAi)* and *par-2(RNAi)* treatments resulted in a reduction in the density asymmetry of the short-lived-population (Figure 5A,S9A) accounting for the centred final spindle-position (Figure S9B). Indeed, the asymmetric distribution of GPR-1/2 is PAR-dependent (Gotta and Ahringer, 2001; Colombo *et al.*, 2003; Gotta *et al.*, 2003; Srinivasan *et al.*, 2003; Tsou *et al.*, 2003; Pécréaux *et al.*, 2006a; Fielmich *et al.*, 2018; Rodriguez-Garcia *et al.*, 2018). Importantly, it did not affect the mitotic progression, suggesting this latter control may be independent of the force-generator density. Indeed, in both depletions, we observed a strong increase in the lifetimes of the short-lived and long-lived populations (Figure 5BC), consistent with pulling force increase along mitosis. However, all cortical regions were equally affected, maintaining the anteroposterior symmetry of the lifetimes in treated and control conditions. It further shows that the force-generator processivity does not encode the pulling-force imbalance. Overall, we suggest that the polarity and mitotic progression controls act independently, respectively through the dynein on-rate (density of active force generators) and dynein off-rate (their processivity). Furthermore, PAR-2 and PAR-3 proteins play an additional role in globally scaling, likely indirectly, microtubule residence times at the cortex.

**Figure 5:**
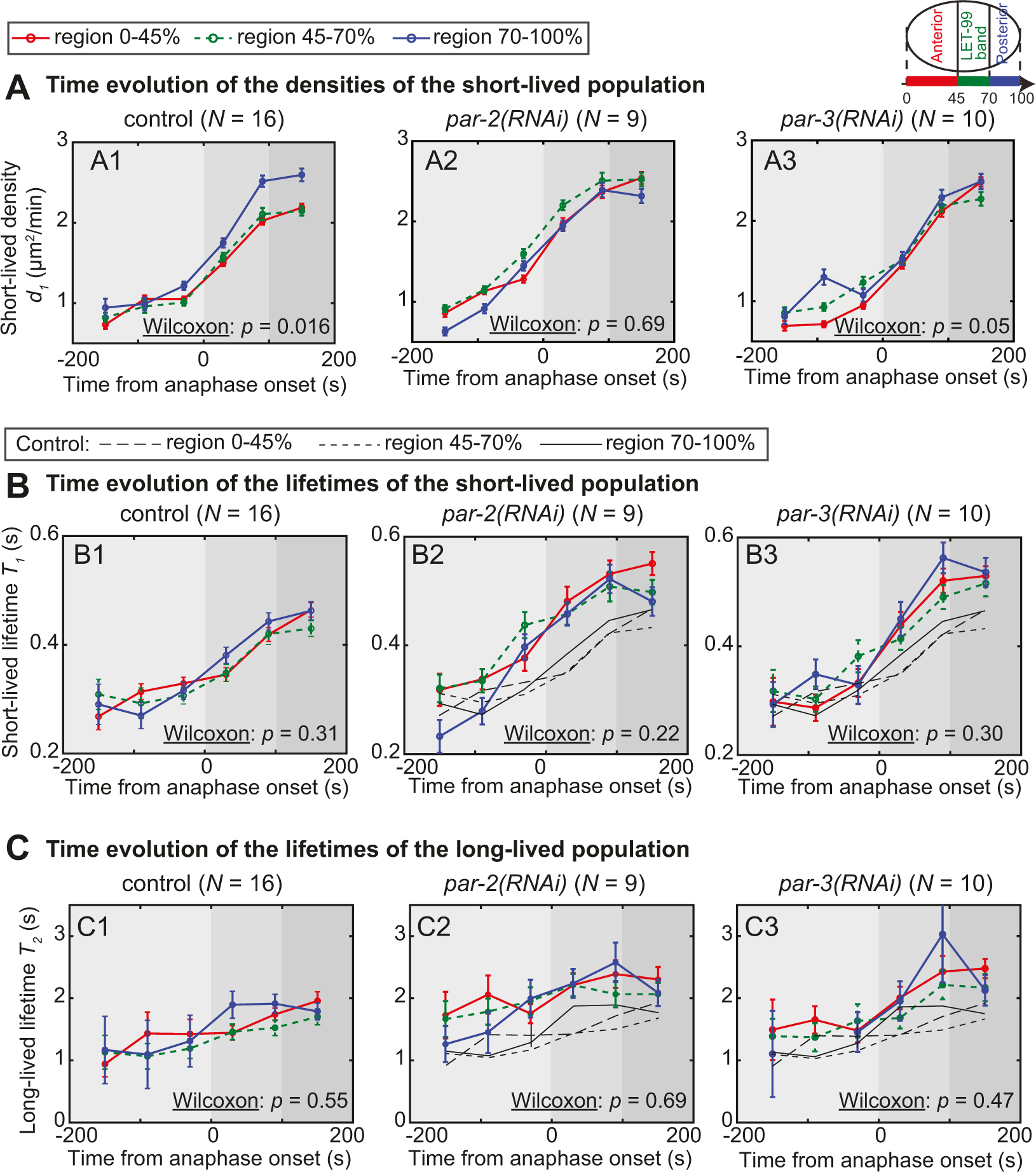
The PAR proteins control the polarisation of the short-lived microtubule density and the residence time of both populations. (**A-C**) Evolution of the DiLiPop parameters during metaphase and anaphase in (red) the anterior region, (green) the lateral LET-99 band and (blue) the posterior-most region: (**A**) short-lived densities, (**B**) short-lived lifetimes, and (**C**) long-lived lifetimes. The three cortical regions are depicted in the schematics at the top right. We analysed, using 60-s time-blocks, *N* = 16 control embryos (left), *N* = 9 *par-2(RNAi)-*treated embryos (middle) and (A3) *N* = 8 *par-3(RNAi)-*treated embryos (right). Standard deviations were computed by bootstrapping (Supp. Text, §1.2.5). We tested a significant difference between the anterior and posterior-most time-series with the Wilcoxon rank test. In (B2), (B3), (C2) and (C3), thin black lines report the corresponding controls. The grey shadings depict, from lighter to darker, the three time-periods, metaphase (the 200 s before anaphase onset), early anaphase (the 100 s after anaphase onset), and late anaphase (from 100s to 200s after anaphase onset).

We next asked whether the microtubules could push against the cortex asymmetrically and displace the spindle posteriorly. Indeed, such a mechanism was proposed and modelled in other organisms (Pavin *et al.*, 2012; Zhao *et al.*, 2012). To test this possibility, we investigated the temporal evolution of the long-lived-population parameters using 60-s time-blocks. We measured symmetric densities until mid-anaphase (Figure S10A1,A3,B1). The lifetimes in anterior and posterior-most regions were quite similar. They were larger anteriorly only in late anaphase (Figure S10A2,A4,B2). Because this asymmetry happened later than the spindle posterior displacement, it suggests that microtubule growing may not contribute to the causative force imbalance.

To gain certainty, we increased the force due to pushing microtubules. EFA-6^PSD^ was reported to negatively regulate both dynein-dependent pulling-force generator and cortical microtubule stability (O’Rourke *et al.*, 2007; O’Rourke *et al.*, 2010). The DiLiPop analysis of *efa-6(RNAi)*-treated embryos showed a modest increase in the short-lived microtubule density (Figure S11A1,B) and a stronger increase in the long-lived one (Figure S11A2,C). In the same condition at the spindle plane, we observed reduced peak-to-peak oscillation-amplitudes for the posterior centrosome (2.33 ± 1.60 µm, *N* = 11) compared to control embryos (5.11 ± 0.90 µm, *p* = 2.5×10^−4^, *N* = 8), as expected (O’Rourke *et al.*, 2010). Importantly, we only observed a slightly increased posterior displacement of the posterior centrosome, however non-significant (Figure S11D). It contrasts with the large increase of the long-lived microtubule density and suggests that microtubule pushing is unlikely to contribute to the posterior displacement. We recently suggested that it maintains the spindle in the cell centre instead (Pécréaux *et al.*, 2016).

We next wonder how independent the positional control is from the polarity one. We recently proposed that the dynein on-rate is decreased by a scarcity of microtubule contacts in the posterior-most region, when the centrosome is far from it during metaphase (Krueger *et al.*, 2010; Bouvrais *et al.*, 2018). Since the spindle is shifted towards the posterior from late-metaphase to the mitosis end, this positional control could contribute to the pulling-force imbalance. Both long-lived and short-lived populations would undergo such a geometrical effect while polarity control affects only the short-lived microtubules. We thus measured the density of long-lived microtubules and observed an asymmetry only in late anaphase (Figure S10A1,B1). At that time, the spindle already migrated posteriorly. Therefore, the positional control may reinforce the posterior displacement lately but not cause it.

To gain certainty about this positional control role, we used again CNSK-1 depletion to alter polarity. Consistently, the time-resolved measurement of short-lived microtubule density shows no significant asymmetry. We noticed a slight anterior enrichment in metaphase (Figure S6A3,B1). Importantly, we measured a global up scaling of the long-lived densities, but no alteration of their spatial distribution in comparison to the control (Figure S6C). Because both populations would be affected equally by a positional control, and since *cnsk-1(RNAi)* altered only the distribution of the short-lived population, the polarity regulation appears sufficient to control the spindle displacement out of the cell centre. We concluded that positional and polarity controls are independent. Again, the positional control can reinforce the asymmetry later in anaphase and may account for the twice-larger posterior force, while the short-lived anterior-to-posterior density ratio is lower (Grill *et al.*, 2001; Grill *et al.*, 2003).

Lastly, to ascertain that the sole asymmetry of dynein density, due to its on-rate, accounts for force imbalance, we asked whether the final position of the spindle correlated with the posterior short-lived-population enrichment. We tested the correlation of the final spindle position along the AP axis and the anterior-to-posterior density ratio of the two populations, during anaphase (Material and Methods). We obtained a more pronounced anti-correlation for the short-lived microtubules (Figures 4C, S10C). Interestingly, the spindle centred position was estimated by linear regression to correspond to a ratio equal to 1.07 for the short-lived population (Figure 4C, dotted grey line) – an almost symmetric distribution. In a broader take, we concluded that the pulling force imbalance is recapitulated by the asymmetric density ratio of the short-lived population. In turn, this density would correspond to the binding rate of dynein to microtubule or its run-initiation.

### The mitotic progression controls the force generator processivity

The cortical pulling force increased during mitosis (Labbé *et al.*, 2004; McCarthy Campbell *et al.*, 2009). In our modelling of pulling force, we attributed it to the increasing processivity of the force generator (Pécréaux *et al.*, 2006a; Bouvrais *et al.*, 2018). The dynein processivity being reflected in the short-lived microtubule lifetime in our assay, the DiLiPop offers an opportunity to validate such a mechanism at the microscopic scale. We measured the temporal evolution of the two lifetimes using 30-s time-blocks but not distinguishing various regions to gain certainty and because we excluded that lifetimes contributed to the force imbalance. We found a steep increase in the short-lived microtubule lifetime during the early anaphase, continued by a shallower one in late anaphase (Figure 6A1). In contrast, the lifetime remained constant during metaphase. Such a variation accounts for the force measured at cell scale. However, we also observed the same increase-pattern for the long-lived population (Figure 6A2) although the variation amplitude is reduced, especially considering relative values. Importantly, both time-series are likely independent during metaphase (Pearson *r* = 0.73, *η*^2^ 2 test *p* = 0.098). It may suggest a specific regulation of the lifetime of the short-lived population, which would superimpose to a general regulation visible on both populations.

**Figure 6:**
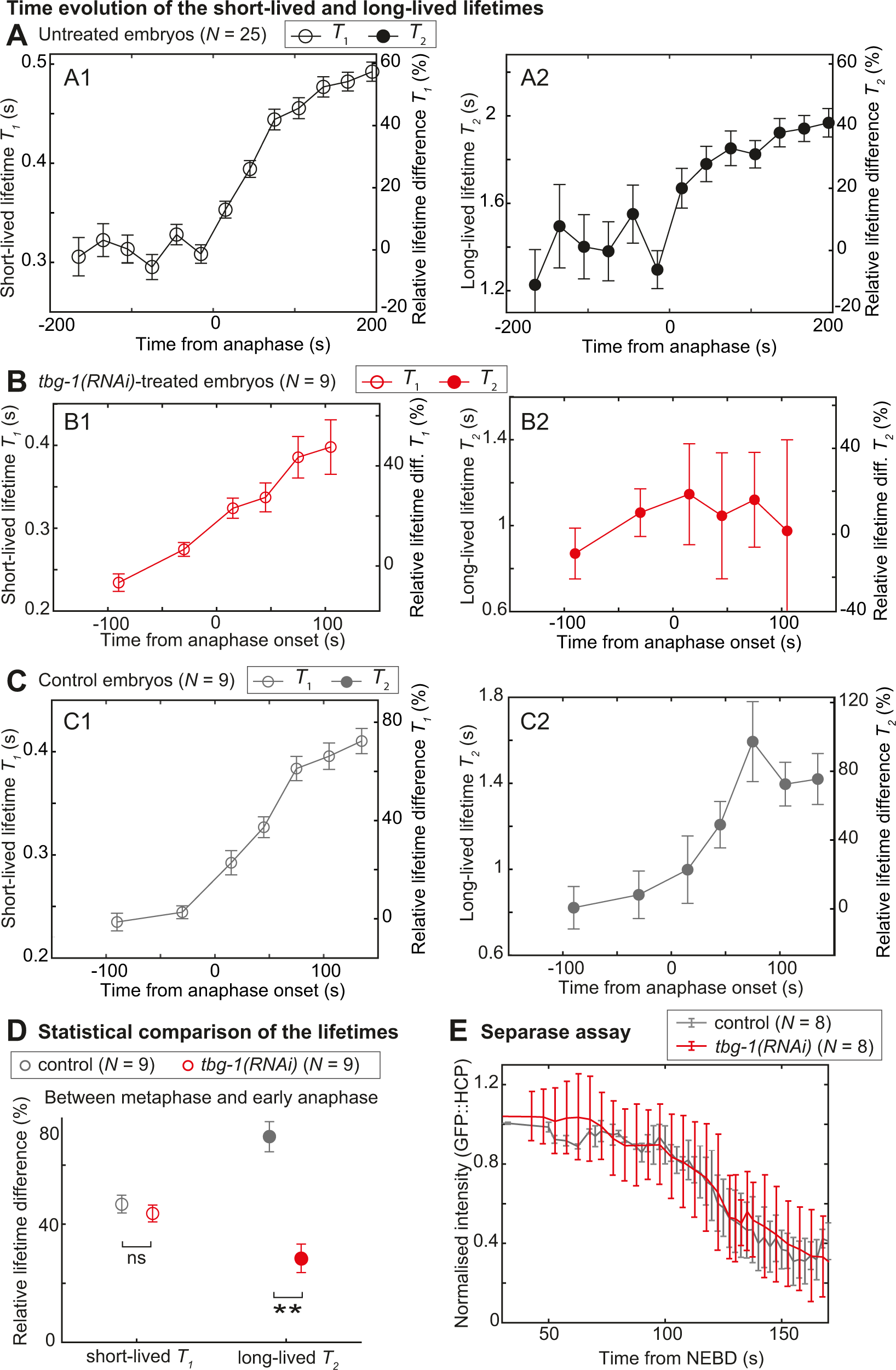
The short-lived population lifetime increases sharply during mitosis, independently of spindle posterior displacement. (**A-C**) Temporal evolutions of the microtubule lifetimes of (A, black) *N* = 25 untreated embryos for (A1) short-lived and (A2) long-lived populations (same data as in Figures 2A, 4AB, S10AB), (B, red) *N* = 9 *tbg-1(RNAi)*-treated embryos, and (C, grey) their control embryos (*N* = 9). We considered a single region encompassing the whole cortex and used 30-s time-blocks for untreated embryos. For *tbg-1(RNAi)-*treated embryos and their controls, we used 60-s time-blocks during metaphase and 30-s time-blocks during anaphase. Y-scale on the right-hand side displays relative lifetime difference from metaphase mean value. Error bars were obtained by bootstrapping (Supp. Text, §1.2.5). The long-lived-lifetime time-series of the control and *tbg-1(RNAi)-*treated embryos were independent (Pearson *r* = 0.35, *η*^2^ test *p* = 0.49), while the short-lived-lifetime ones were correlated (*r* = 0.98, *p* = 7×10^−4^). (**D**) Differences in short- and long-lived lifetimes between metaphase and early anaphase (from 0 s to 100 s from anaphase onset), normalised by their respective metaphase lifetimes for (red) *N* = 9 *tbg-1(RNAi)-*treated embryos and (grey) *N* = 9 control embryos. Stars indicate significant differences (Student’s *t*-test). (**E**) Mean chromosomal GFP fluorescence of the separase sensor over time, for (red) *N* = 8 *tbg-1(RNAi)-*treated embryos and (grey) *N* = 8 control embryos. Error bars indicate standard deviations.

We sought a condition perturbing the lifetime of one of the two populations, to test such a regulation difference. We depleted the microtubule rescue factor CLS-2^CLASP^. It is expected to affect astral microtubule quite independently of cortical pulling force generators (Srayko *et al.*, 2005; Espiritu *et al.*, 2012). We kept the *cls-2(RNAi)* treatment hypomorphic to ensure functional spindle and central spindle (Cheeseman *et al.*, 2005; Maton *et al.*, 2015). We observed a different evolution of short- and long-lived microtubule lifetimes during late anaphase (Figure S12A). We also found a significant reduction in the long-lived microtubule lifetime after mid-anaphase, while the short-lived lifetime was only slightly downscaled (Figure S12C). Furthermore, while the short-lived times-series of *cls-2(RNAi)-*treated and control embryos were correlated, the long-lived ones were mildly independent (Figure S12AB). It suggests that CLS-2 depletion affected mostly the long-lived population. Under the same condition and at the spindle plane, we measured a faster spindle elongation equal to 0.517 ± 0.287 µm/s upon *cls-2(RNAi)* (*p* = 9.8×10^−4^, *N* = 10) compared to 0.083 ± 0.027 µm/s for the control embryos (*N = 7*), confirming the penetrance of the RNAi treatment (Espiritu *et al.*, 2012). Because CLS-2 is a rescue factor, likely, it is relevant that microtubules involved in generating pushing-force are especially affected. In all case, it suggests that the lifetimes may be differentially regulated between both populations.

We next sought an alteration of the spindle position not impairing pulling-force regulation. Indeed, it could reveal whether the two lifetimes are separately regulated. We set to reduce the spindle to a single centrosome (or two centrosomes not clearly separated) performing a *tbg-1(RNAi)* treatment (Motegi *et al.*, 2006). We measured a lifetime of the long-lived population significantly decreased in anaphase compared to the control one, while the short-lived lifetime is only mildly affected (Figure 6B-D). Consistently, the short-lived and long-lived microtubule-lifetime time-series are non-correlated (Pearson *r* = 0.45, *χ*^2^ test *p* = 0.36). It suggests that the short-lived population lifetime is mostly not affected by the centrosome position. We measured a decreased density for both populations during anaphase (Figure S12E) due to the positional control, as expected (Bouvrais *et al.*, 2018). As intended, the posterior displacement was reduced during anaphase (Figure S12D). These observations suggest that another mechanism may control the short-lived population, on top of the global regulation of astral microtubules.

We asked whether the above changes in lifetime evolution could result from an altered regulation of microtubule dynamics or of dynein processivity, due to modified cell cycle progression and particularly at anaphase onset (Srayko *et al.*, 2005; McCarthy Campbell *et al.*, 2009). To do so, we used the separase activity assay (Kim *et al.*, 2015). We performed the same treatments in a strain labelled mCh::H2B and GFP::sensor, the sensor being a readout of separase activity (Material and Methods). We measured fluorescent signal at the chromosomes from NEBD to mid-anaphase. We observed a decrease in GFP fluorescent signal at about 100 s from NEBD for both the control and *tbg-1(RNAi)-*treated embryos (Figure 6E). It confirmed that the separase was activated similarly in the two conditions. It suggested a normal temporal control of the dynein processivity and microtubule dynamics upon *tbg-1(RNAi)*. These results agreed with the strong correlation between the short-lived-lifetime time-series of *tbg-1(RNAi)*-treated embryos and their controls (Pearson *r* = 0.98, *χ*^2^ test *p* = 7×10^−4^). Overall, while a general regulation of the microtubule dynamics exists, we suggest that the short-lived microtubule lifetime increases beyond that regulation. It is consistent with an increasing processivity that causes force build-up (Labbé *et al.*, 2004; Pécréaux *et al.*, 2006a) and accounts for the mitotic-progression control of the cortical pulling force.

### The polymerising microtubules contribute to maintaining the spindle in the cell centre

We recently proposed, from cell scale measurements, that the spindle is maintained in the cell centre during metaphase by microtubules pushing against the cortex (Garzon-Coral et al., 2016; Pécréaux et al., 2016). We monitored the long-lived population, which reveals microtubules pushing against the cortex, to test this hypothesis at the microscopic scale. We varied the long-lived microtubule density by targeting either MAPs (Srayko *et al.*, 2005) or polarity proteins (Labbé *et al.*, 2003; Severson and Bowerman, 2003). We used the stability of the metaphasic spindle in the cell centre (Pécréaux et al., 2016), measured through the diffusion coefficient of the spindle position along the transverse axis *D*Sy computed from images taken at the spindle plane (Berg-Sørensen and Flyvbjerg, 2004; Nørrelykke and Flyvbjerg, 2010). The smaller this value, the better the centring stability. We found an anti-correlation between this measurement and the density of long-lived microtubules (Figure 7A) but not with the short-lived density (Figure 7B). These direct observations of force-generating events suggest that microtubules pushing – rather than pulling – contribute to maintaining the spindle in the cell centre. Furthermore, because the microtubule density at the cortex impacted the centring, these results are not consistent with the cytoplasmic pulling hypothesis.

**Figure 7:**
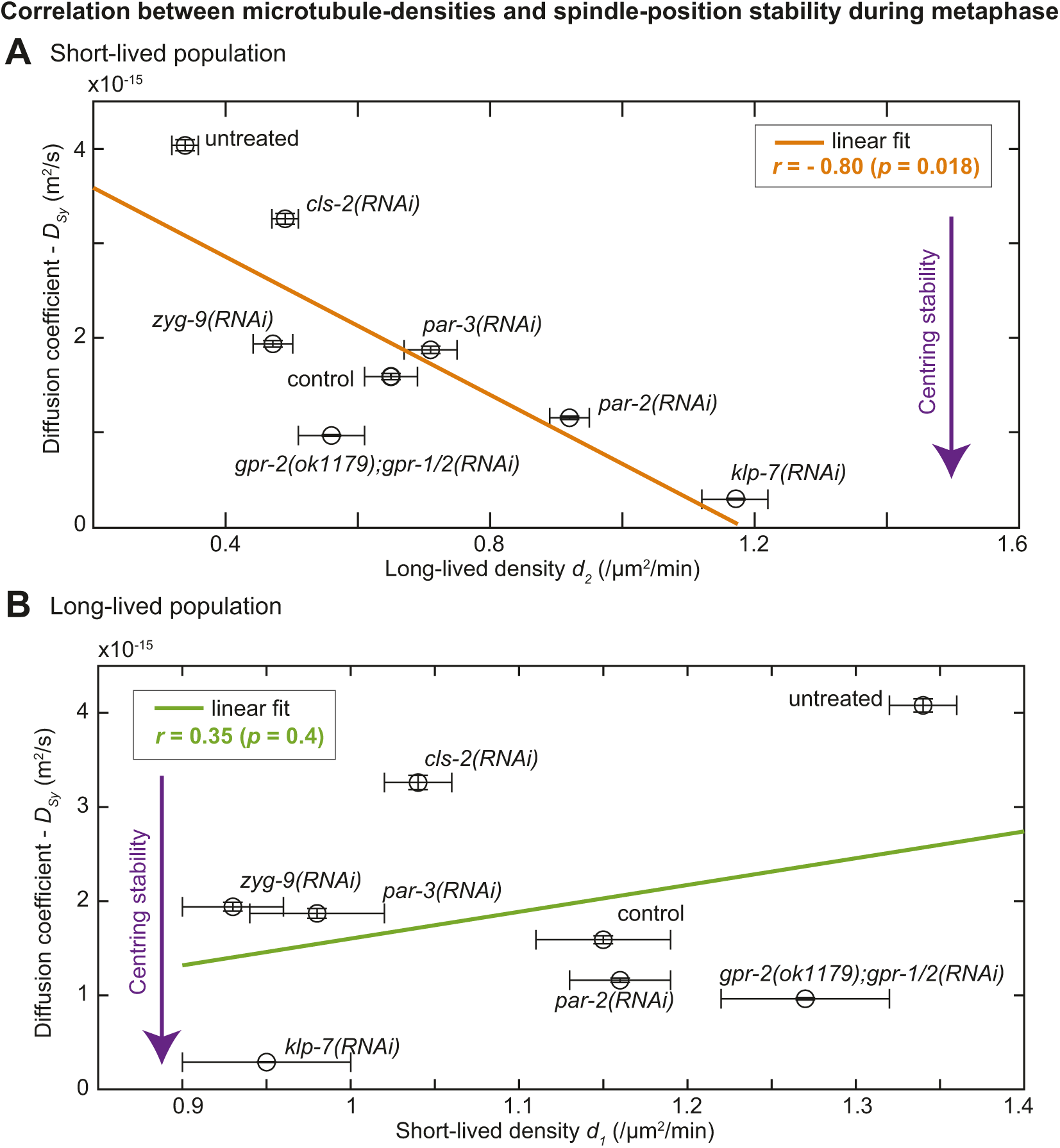
The long-lived microtubules, reflecting pushing forces, contribute to maintaining the spindle at the cell centre. Diffusion coefficient of the spindle position along the transverse axis, *D*_*Sy*_, characterizing the centring stability and based on imaging at the spindle plane, plotted against the density of the (**A**) long-lived and (**B**) short-lived microtubule density during metaphase obtained by DiLiPop analysis of images at the cortex. The orange and green lines depict the Pearson correlations, respectively, for the long-lived and short-lived populations. We varied the pulling and pushing forces by using *klp-7(RNAi)* (*N* = 8 at the cortex and *N* = 9 at the spindle plane, written as 8/9 for the following conditions), *zyg-9(RNAi)* (*N* = 13/8), *cls-2(RNAi)* (*N* = 11/9), *par-2(RNAi)* (*N* = 9/9), *par-3(RNAi)* (*N* = 10/6), *gpr-2(ok);gpr-1/2(RNAi)* (*N* = 8/8), control embryos (*N* = 8/10) and untreated embryos (*N* = 10/10).

## DISCUSSION

Through an advanced and careful analysis of microtubule-contact dynamics at the cortex, we monitored the distribution of two microtubule populations distinct by their residence times. Our measured lifetimes, 0.4 s and 1.8 s, are short compared to previously published values, which range between 1 s and 15 s (Labbé *et al.*, 2003; Kozlowski *et al.*, 2007; O’Rourke *et al.*, 2010; Lacroix *et al.*, 2016; Schmidt *et al.*, 2017; Sugioka *et al.*, 2018). Most of these previous measurements were obtained labelling only the growing microtubules through EB proteins. Interestingly, approaches with higher frame rates, consistent with microtubule growth and shrinkage rates, provide smaller residence times, close to the values found here. It may suggest that high frame rates are needed to resolve the exceptionally fast dynamics in the nematode compare to other organisms (Chaaban *et al.*, 2018). Beyond measuring the residence time, we aimed to understand the regulation of the forces positioning the spindle by analysing the statistics of individual events. Importantly to ensure the sampling is representative, we estimate that DiLiPop recovers about 66 % of the microtubule contacts at the cortex, based on electron micrographs (Redemann *et al.*, 2017). Overall, the DiLiPop being accurate and representative enabled us to decipher and quantitatively understand the complex force regulations that conduct the spindle choreography.

We interpreted dynamically distinct populations as microtubule pushing and pulling events, respectively corresponding to long- and short-lived cortical contacts. Interestingly, upon perturbing by RNAi either microtubule dynamics regulators or the cortical force-generating complex, the population proportions changes but not the total contact count. Such an observation suggests that the belonging to a population for a microtubule is a dynamical choice. It depends likely whether the microtubule meets or not a (rare) trimeric force-generating complex at the cortex (Grill and Hyman, 2005; Pécréaux *et al.*, 2006a; Park and Rose, 2008; Riche *et al.*, 2013; Bouvrais *et al.*, 2018). In contrast, this finding is poorly consistent with cortical residence-time differing due to the microtubule aging or post-translational modification (PTM) (Srayko *et al.*, 2005; Portran *et al.*, 2017; Lacroix *et al.*, 2018; Schaedel *et al.*, 2019). Furthermore, the labelling of *α*- or *β*-tubulin only mildly scaled the lifetimes, likely because of dye-brightness difference, while the proportions were preserved. This independence from labelled tubulin paralogs is hardly consistent with PTM regulating the microtubule lifetime at the cortex. Overall, DiLiPop offers a dynamical read-out of the distribution of force-generating events in space and time.

### The short-lived population may also include stalled microtubules

We surprisingly measured the anterior-to-posterior density ratio of the short-lived microtubules to be about 0.88, neatly above 0.5. It contrasts with the accepted view of twice more active force-generators at the posterior cortex compared to the anterior one (Grill *et al.*, 2003). Interestingly, analysing the dynein dynamics at the cortex led to a similar ratio for the dynein molecules involved in pulling (Rodriguez-Garcia *et al.*, 2018). These non-pulling events could correspond to stalled microtubule-ends/dyneins. Indeed, *in vitro* and *in vivo* studies showed that anchored dynein could serve as microtubule plus-end tether (Dujardin and Vallee, 2002; Hendricks *et al.*, 2012; Laan *et al.*, 2012b; Perlson *et al.*, 2013; Yogev *et al.*, 2017; Bouvrais *et al.*, 2018). Consistently, the number of short-lived microtubules contacting the cortex is larger than the expected values of 10-100 per cortex half (Grill *et al.*, 2003; Redemann *et al.*, 2010). During late anaphase, we measured about 40 short-lived microtubules contacting the visible cortex each second, extrapolated to about 120 per half cortex. It can reveal a mechanism regulating dynein run initiation from a stalled state to bound to a microtubule (Laan *et al.*, 2012a; Jha *et al.*, 2017).

### The pushing force maintains the spindle in the cell centre during metaphase

The final position of the spindle results from the balance of centring and pulling forces (Pécréaux *et al.*, 2006a; McNally, 2013; Bouvrais *et al.*, 2018). Our approach allowed us to investigate how the spindle is maintained in the cell centre during metaphase at the scale of a single microtubule. Indeed, we recently proposed that the microtubule pushing against the cortex could account for the extraordinary accuracy of this positioning (Pécréaux *et al.*, 2016). Consistently, during metaphase, we observed that the density of long-lived microtubules correlates with centring stability. In contrast, the short-lived density measurements appear poorly correlated with the centring stability. Furthermore, this population displays a reduced density during metaphase compared to anaphase. It is consistent with the pulling force contributing to de-centring (Dogterom *et al.*, 2005; Grill and Hyman, 2005; Kozlowski *et al.*, 2007; Zhu *et al.*, 2010; Garzon-Coral *et al.*, 2016; Pécréaux *et al.*, 2016). Overall, single microtubule-contact analysis suggests that the centring mechanism is due to microtubules pushing against the cortex.

Recently, the APR-1/APC complex was suggested to decrease the cortical forces anteriorly as it measured a reduced lifetime at the anterior cortex (Sugioka *et al.*, 2018). This study differs by the method to distinguish populations. Consequently, our results contrast and we did not observe an increased density or lifetime of the long-lived population anteriorly during anaphase. It suggests that the centring force does not contribute to the posterior displacement. Our study also supports successive dominance of pushing and pulling along time (Ahringer, 2003; Pécréaux *et al.*, 2006a; Garzon-Coral *et al.*, 2016; Bouvrais *et al.*, 2018). During metaphase, the pulling force plateaus. It results in only a slow posterior displacement but lets the centring forces dominate along the transverse axis (Grill *et al.*, 2001; Ahringer, 2003; Garzon-Coral *et al.*, 2016; Pécréaux *et al.*, 2016). In anaphase, the pulling reinforces. We observed that the short-lived-microtubule lifetime undergoes a more pronounced increase compared to the long-lived lifetime. This regulation through intensifying the pulling/displacement forces contrasts with recent findings in the sea urchin zygote whereby a reduction of the centring forces accounts for the de-centration after the maintenance in cell centre (Sallé *et al.*, 2018). In the nematode zygote, pushing force barely superimposes to the pulling one without contributing to the asymmetric positioning of the spindle (Grill and Hyman, 2005; Pécréaux *et al.*, 2006a).

### The cortical pulling control is threefold, by mitotic progression, polarity and the spindle position

We recently proposed that a second regulation of the pulling force, by position of the centrosomes, superimposed to the mitotic progression control reflected in the processivity of the force generators (Figure 8, respectively left and right blocks) (Pécréaux *et al.*, 2006a; Bouvrais *et al.*, 2018). The DiLiPop sheds light on the interplay of these controls with the polarity one reflected in the asymmetry of dynein on-rate (Figure 8, middle block) (Rodriguez-Garcia *et al.*, 2018). Beyond confirming that the dynein detachment rate does not encode the polarity (Figure 8, mixed pink/purple boxes) (Rodriguez-Garcia *et al.*, 2018), we broadly found no other cause of force imbalance. Importantly, we observed that this asymmetry is set early in the division and is scaled up by the global and symmetric increase in processivity viewed through short-lived microtubule lifetime (Figure 8, purple boxes). Such a mitotic progression control is consistent with the previous measurements at the cell-scale (Labbé *et al.*, 2004; Pécréaux *et al.*, 2006a; McCarthy Campbell *et al.*, 2009). This scaling is likely not gradual. Indeed, we observed a steep increase in the cortical residence time at anaphase onset.

**Figure 8:**
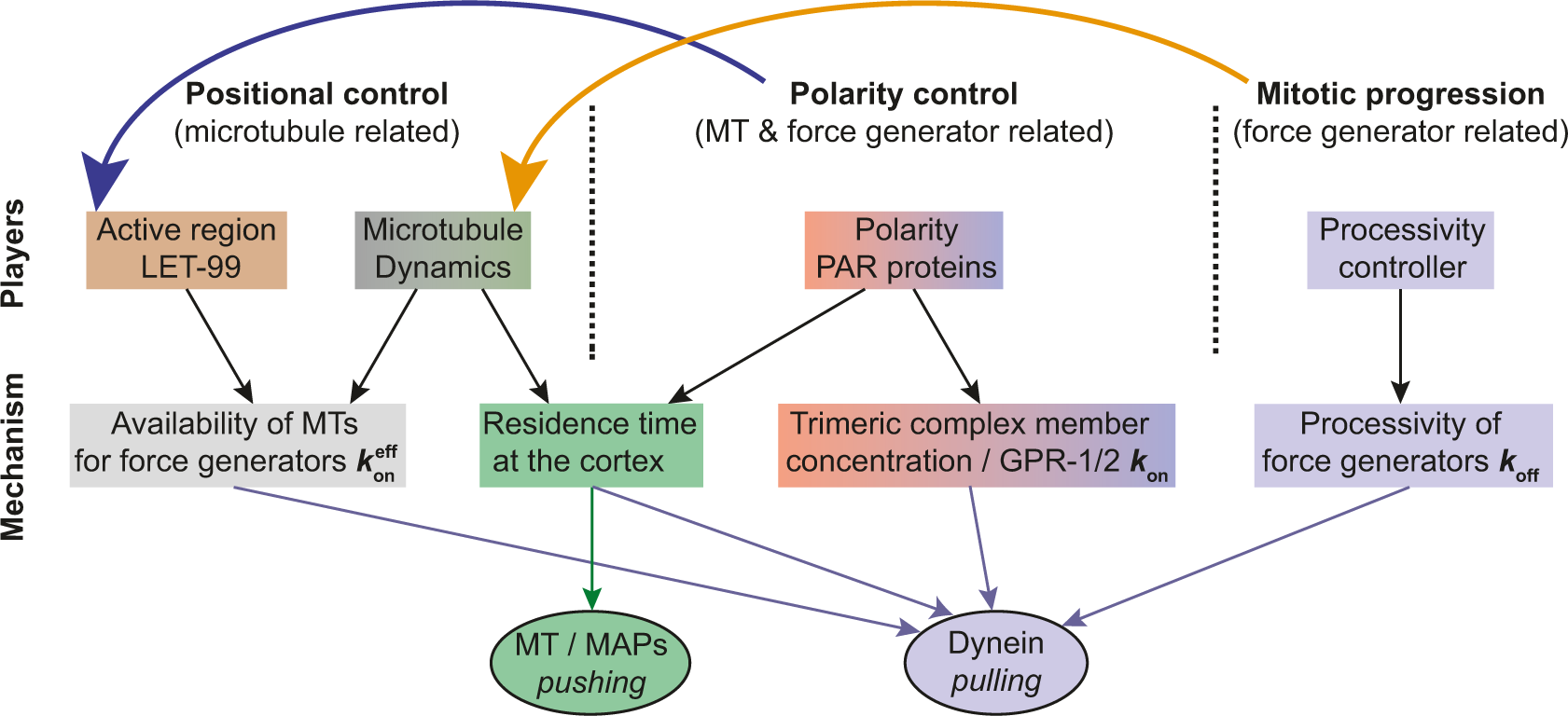
The cortical pulling control is threefold, by mitotic progression, polarity and the spindle position. Schematics of the regulation of force-positioning the spindle with the players (top row), and the quantity regulated (middle row). Grey and brown colours correspond to the positional control involving astral microtubule (MT) dynamics and the active region created by LET-99 band. Purple colour depicts the time control through force generator processivity. Pink/purple colours correspond to the polarity control involving the distribution of the force generators. The latter control also participates in setting the microtubule residence time at the cortex (green).

Finally, the DiLiPop suggests that the positional control only reinforces the anteroposterior imbalance of cortical pulling forces in late anaphase (Figure 8, grey box). Consistently, the long-lived microtubule density becomes slightly asymmetric only in late anaphase (Figure 8, brown box) (Riche *et al.*, 2013; Bouvrais *et al.*, 2018). While not polarized in early mitosis, this control is affected by PAR-2/PAR-3 proteins, which decrease microtubule lifetimes of both populations (Figure 8, green box). The positional control contributes to force imbalance in late anaphase, and this mechanism depends on the posterior-most region created by the LET-99 protein. Establishing this protein domain is under the control of the polarity (Figure 8, top blue arrow) (Wu and Rose, 2007; Krueger *et al.*, 2010; Wu *et al.*, 2017b). Both cross talks create a loose link between the polarity and the positional control. On the side of the mitotic progression, the cell cycle controls the number of nucleated microtubules, known to increase at anaphase (Srayko *et al.*, 2005). It increases the microtubule density of both populations, symmetrically, connecting mitotic and positional controls (Figure 8, top orange arrow). However, such a link is loose, and the controls remain mostly independent (Bouvrais *et al.*, 2018).

### Conclusion

Overall, we propose that the pulling forces are under three independent controls: polarity reflected as the force-generator on-rate due to an asymmetric distribution of GPR-1/2; mitotic progression, corresponding to the processivity of the force generators; positional control, due to the availability of the microtubules at the cortex. The centring mechanism is due to microtubules pushing against the cortex and barely superimposes to the pulling forces. Beyond these findings, this work exemplifies the interest of combining investigations at two scales. In particular, it offers the unparalleled ability to view the individual pulling and pushing force-generating events. We foresee that this novel approach will find applications beyond cell division.

## MATERIALS AND METHODS

### Culturing C. elegans

*C. elegans* nematodes were cultured as described in (Brenner, 1974), and dissected to obtain embryos. The strains were maintained at 25°C and imaged at 23°C. The strains were handled on nematode medium plates and fed with OP50 bacteria.

### Strains of C. elegans and C. briggsae used

*elegans* TH65 YFP::TBA-2 (α-tubulin) strain (Srayko *et al.*, 2005) having a fluorescent labelling of the whole microtubule (MT) was used for the DiLiPop assay as well as *C. elegans* AZ244 GFP::TBB-2 (β-tubulin) strain (Praitis *et al.*, 2001) and *C. briggsae* ANA020 GFP::TBB (β-tubulin) strain. TH65 strain was also the standard for the “centrosome-tracking” assay used to validate the penetrance of RNAi treatments. TH66 EBP-2::GFP strain (Srayko *et al.*, 2005) that displays a labelling of microtubule plus-ends was used for comparison of its effects on microtubule dynamics. The JEP18 *gpr-2(ok1179)* strain was used to target GPR-1/2 protein though a mutation. The OD2207 strain, expressing HIS-58 fused to mCherry and a sensor composed of GFP fused to the CPAR-1 N-tail placed in front of the histone fold domain (HFD) of HCP-3, was used for the separase assay (Kim *et al.*, 2015).

### Gene inactivation through protein depletion by RNAi feeding

RNA interference (RNAi) experiments were performed by feeding using the Ahringer-Source BioScience library (Kamath and Ahringer, 2003), except for GOA-1;GPA-16 depletion, whose clone was kindly given by Prof P. Gönczy. The feedings were performed at 25°C for various durations according to the experimental goals. The treatment lasted 24h for *lin-5, goa-1*;*gpa-16* and *klp-7* genes. When we aimed for stronger phenotypes (e.g. symmetric divisions), we used duration of 48h (*cls-2, par-2, par-3* and *gpr-1/2*). The duration was reduced to 4h and 6-10h when targeting *zyg-9, efa-6* and *cnsk-1*, respectively. The control embryos for the RNAi experiments were fed with bacteria carrying the empty plasmid L4440. We did not notice any phenotype suggesting that the meiosis was impaired during these various treatments.

### Preparation of the embryos for imaging

Embryos were dissected in M9 buffer and mounted on a pad (2% w/v agarose, 0.6% w/v NaCl, 4% w/v sucrose) between a slide and a coverslip. Depending on the assay (landing or centrosome-tracking ones), embryos were observed using different microscopic setups. To confirm the absence of phototoxicity and photodamage, we checked for normal rates of subsequent divisions (D.L. Riddle, 1997; Tinevez, 2012). Fluorescent lines were imaged at 23°C.

### Imaging of microtubule contacts at the cortex

We imaged *C. elegans* one-cell embryos at the cortex plane in contact with the glass slide, viewing from the nuclear envelope breakdown (NEBD) until late anaphase. We used a Leica DMi8 spinning disk microscope with Adaptive Focus Control (AFC) and a HCX Plan Apo 100x/1.4 NA oil objective. Illumination was performed using a laser with emission wavelength of 488 nm and we used GFP/FITC 4 nm band pass excitation filter and a Quad Dichroic emission filter. To account for the fast microtubule dynamics at the cortex, images were acquired at an exposure time of 100 ms (10 Hz) using an ultra-sensitive Roper Evolve EMCCD camera that was controlled by the Inscoper device (Combo Microtech). During the experiments, the embryos were kept at 23°C. To image embryos at the cortex, we typically moved the focus to 12 to 15 µm below the spindle plane. Images were then stored using Omero software (Li *et al.*, 2016).

### Spindle pole imaging

Embryos were observed at the midplane using a Zeiss Axio Imager upright microscope modified for long-term time-lapse. First, an extra anti-heat filter was added to the mercury lamp light path. Secondly, to decrease the bleaching and obtain optimal excitation, we used an enhanced transmission 12 nm band pass excitation filter centred on 485 nm (AHF analysentechnik). We used a 100x/1.45 NA Oil plan-Apo objective. Images were acquired with an Andor iXon3 EMCCD 512×512 camera at 33 frames per second and using their Solis software. Images were then stored using Omero software (Li *et al.*, 2016).

### Centrosome-tracking assay

The tracking of labelled centrosomes and analysis of trajectories were performed by a custom tracking software (Pécréaux *et al.*, 2006a) and developed using Matlab (The MathWorks). Tracking of −20°C methanol-fixed γ-tubulin labelled embryos indicated accuracy to 10 nm. Embryo orientations and centres were obtained by cross-correlation of embryo background cytoplasmic fluorescence with artificial binary images mimicking embryos, or by contour detection of the cytoplasmic membrane using background fluorescence of cytoplasmic TBG-1::GFP with the help of an active contour algorithm (Pécréaux *et al.*, 2006b). The results were averaged over all of the replicas for each condition.

### Simulation of microscopy images

To validate the image-processing pipeline (Figure S2AB), we built fluorescence images of known dynamics, which mimic our cortical images using the algorithm developed by (Costantino *et al.*, 2005) that we adapted to our needs as previously done (Bouvrais *et al.*, 2018). In further details, we simulated stochastic trajectories of particles that displayed a limited random motion characterized by the diffusion coefficient *D*. We sampled the duration of the tracks from an exponential distribution. We encoded the fluorescence intensity through the quantum yield parameter (Qyield). After plotting the instantaneous positions, we mimicked (1) the effect of the point-spread function (PSF) in fluorescence microscopy by applying a Gaussian filter, and (2) the background noise by adding at each pixel a sampling of a Gaussian distribution. Details of the parameters used for simulation can be found in Table S3.

### Separase sensor assay

To check whether the cell cycle was unaffected by the *tbg-1(RNAi)* treatment, we performed the separase sensor assay introduced in (Kim *et al.*, 2015) using the strain OD2207 (Figure 6E). We acquired 5 x 2 um z-stacks every 2.5 s from NEBD to post chromatid separation. To quantify fluorescence, we used ImageJ (Fiji) and followed the image-processing protocol described in (Kim *et al.*, 2015).

### Statistics

For classic statistical analyses, averaged values of two conditions were compared using the two-tailed Student’s *t*-test with correction for unequal variance except where otherwise stated. The Wilcoxon signed rank test was used to assess whether two time-series of DiLiPop densities/lifetimes were significantly different all along. The Pearson *χ*^2^ test was used to indicate whether two sets of data were correlated or independent. For the sake of simplicity, we recorded confidence levels using diamond or stars (◊, *p* ≤ 0.05; *, *p* ≤ 0.01; **, *p* ≤ 0.001; ***, *p* ≤ 0.0001; ****, *p* ≤ 0.00001) and ns (non-significant, *p* > 0.05; sometimes omitted to save room). We abbreviated standard deviation by SD, standard error by s.e., and standard error of the mean by s.e.m.

### Data and image processing

All data analysis was developed using Matlab (The MathWorks).

## ACKNOWLEDGMENTS

The bacterial clone of GPA-16;GOA-1 was a kind gift from Prof P. Gönczy. We thank Dr. Gregoire Michaux for the feeding clone library and technical support. We also thank Drs. Giulia Bertolin, Aurélien Bidaud-Meynard, Sébastien Huet, Benjamin Mercat, Grégoire Michaux, Anne Pacquelet, Xavier Pinson, and Marc Tramier for discussions about the project. Some strains were provided by the Caenorhabditis Genetics Center (CGC), which is funded by National Institutes of Health Office of Research Infrastructure Programs (P40 OD010440; University of Minnesota).

JP was supported by a Centre National de la Recherche Scientifique (CNRS) ATIP starting grant and La Ligue nationale contre le cancer. We also acknowledge Plan Cancer grant BIO2013-02, COST EU action BM1408 (GENiE) and La Ligue contre le cancer (comités d’Ille-et-Vilaine et du Maine-et-Loire). Microscopy imaging was performed at the Microscopy Rennes Imaging Center, UMS 3480 CNRS/US 18 INSERM/University of Rennes 1. Spinning disk microscope was co-funded by the CNRS, Rennes Métropole and Région Bretagne (AniDyn-MTgrant). DF’s postdoctoral fellowship was funded by Région Bretagne (pRISM grant). HB’s postdoctoral fellowship was funded by the European Molecular Biology Organization. TP was supported by the France-BioImaging infrastructure (ANR-10-INBS-04).

