## Supplemental material for "The coordination of spindle-positioning forces during the asymmetric division of the *C. elegans* zygote is revealed by distinct microtubule dynamics at the cortex"

**SUPPLEMENTAL FIGURES, TABLES and MOVIE** for the paper entitled "**The coordination of spindle-positioning forces during the asymmetric division of the *C. elegans* zygote is revealed by distinct microtubule dynamics at the cortex.**"

**Supplemental Figure S1**: The DiLiPop assay includes estimates of the model-parameter accuracy.

**Supplemental Figure S2**: Two dynamically distinct populations of microtubules reside at the cortex in tubulin-labelled embryos.

**Supplemental Figure S3**: *In silico* investigations reveal the statistical power required to resolve three microtubule populations.

**Supplemental Figure S4**: Generating the DiLiPop density maps enables to depict population variations in time and space.

**Supplemental Figure S5**: *In silico* investigations estimate the requirements for the DiLiPop-statistical analysis to resolve two populations.

**Supplemental Figure S6**: The asymmetry of force-generator density correlates with the pulling-force-generator distribution.

**Supplemental Figure S7**: The asymmetry of the short-lived microtubule density reflects the posterior enrichment of the trimeric force-generating complex.

**Supplemental Figure S8**: Depleting members of the trimeric force-generating complex does not affect the time evolution of the short-lived microtubule lifetimes.

**Supplemental Figure S9**: The asymmetry of force-generator density depends on PAR polarity proteins.

**Supplemental Figure S10**: The long-lived microtubule densities are symmetric until mid-anaphase.

**Supplemental Figure S11**: The pushing related microtubule population does not contribute to the spindle posterior-displacement.

**Supplemental Figure S12**: The long-lived microtubule lifetime depends on CLS-2$^{\text{CLASP}}$.

**Table S1**: Parameters used for KUT-pipeline analysis.

**Table S2**: Typical values of the Bayesian Inference Criterion used to select the best model.

**Table S3**: Parameters used to generate images *in silico*.

**Table S4**: Parameters used for NAM-pipeline processing.

**Movie S1**: Fluorescent spots of microtubules contacting the cortex of a *C. elegans* embryo.

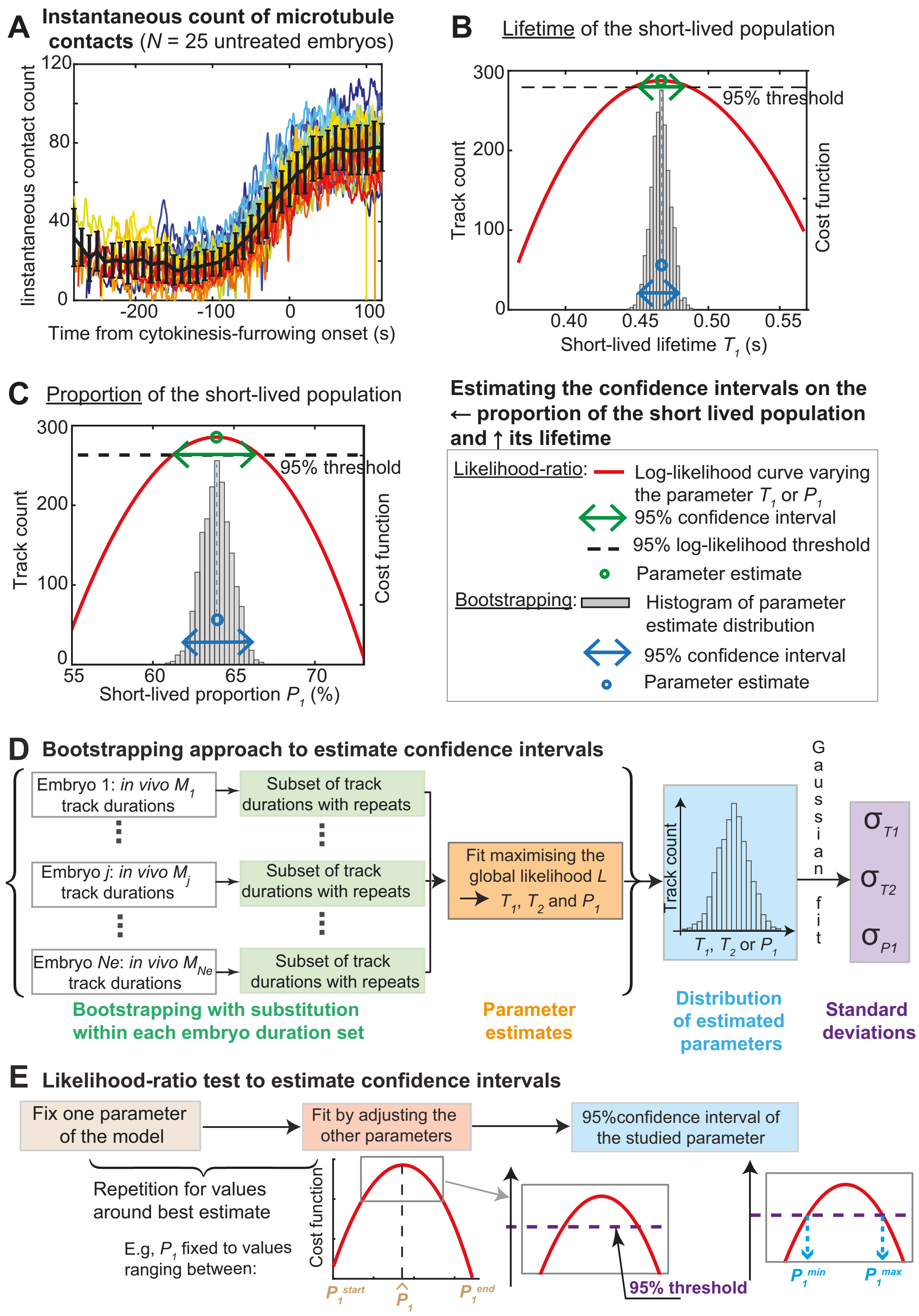


**Supplemental Figure S1: The DiLiPop assay includes estimates of the model-parameter accuracy.**

(**A**) Count of instantaneous microtubule-contacts, i.e. in any single frame, for $N = 25$ untreated embryos (Suppl. Text 1.1.2). Thin coloured lines correspond to individual embryos, while the thick black one to the average. (**B-C**) Exemplar estimating of the confidence intervals on (B) the proportion of the short-lived population $P_1$ and (C) the lifetime of the short-lived population $T_1$ corresponding to the data presented at Figure 1EF (Supp. Text, §1.2.5). The grey bars show the histogram of the parameter estimates obtained by bootstrapping. The blue circle corresponds to its average and the blue arrow to the 95% confidence interval. In the alternative approach using the likelihood ratio, the red curve depicts the cost function (log-likelihood), the green circle its maximum (best estimate), and the green arrow its 95% confidence interval obtained by $\chi^2$ modelling the likelihood (dashed black line). (**D**) (Orange) The microtubule-contract distributions obtained by bootstrapping (Supp. Text, §1.2.5) were fitted to obtain the corresponding optimal parameters. These two steps were repeated typically as many times as the average number of tracks per embryo. (Blue) It resulted in a distribution of parameter estimates. (Purple) By fitting a Gaussian, we extracted a mean and a standard deviation and computed the parameter estimate and its 95% confidence interval. (**E**) In the likelihood-profile approach, (brown shading) we fixed a parameter of the model, and (red shading) we computed the likelihood as a function of this given parameter, other parameters being optimized. (Blue shading) We modelled the resulting cost function with a $\chi^2$ law and determined the corresponding 95% confidence interval (purple dashed line, blue arrows).

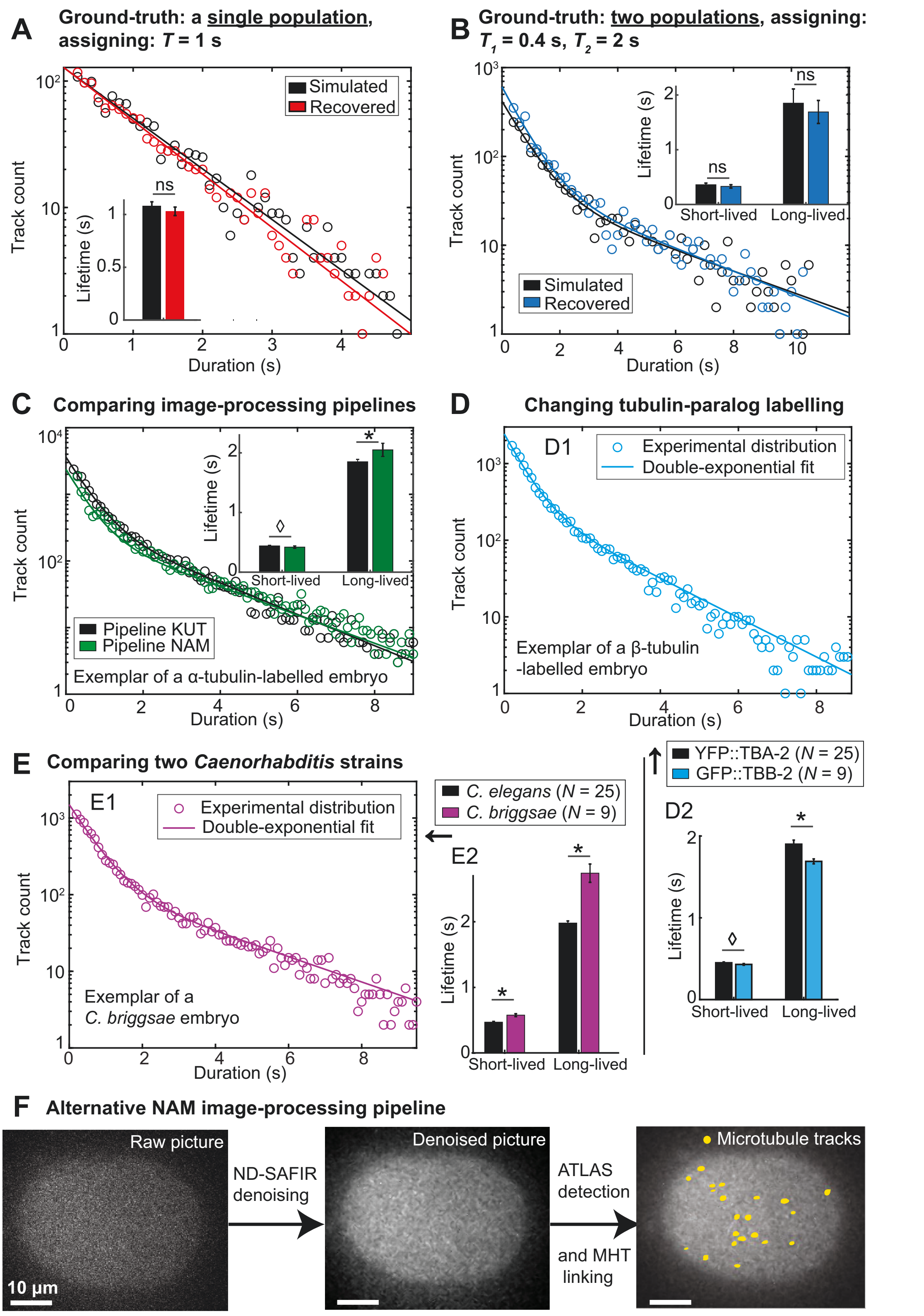


**Supplemental Figure S2: Two dynamically distinct populations of microtubules reside at the cortex in tubulin-labelled embryos.**

We validated the analysis of microtubule dynamics using *in silico* images (Material and Methods). (**A**) (Black circles) Particle lifetime-distribution for a single behaviour, whose lifetime equals to 1 s, was computed and analysed using the KUT-pipeline to get (red circles) the recovered-track duration distribution (parameters in Table S1). Both were best fitted by a mono-exponential, with optimal lifetime shown in inset. (**B**) We performed a similar approach with two populations of lifetimes of 0.4 and 2 s. (**C**) Comparing two different image-processing pipelines using untreated α-tubulin-labelled embryos ($N$ = 20) (parameters in Table S1 and S4) (Supp. Text §1.1.3). The obtained experimental distributions using (black) KUT pipeline and (green) NAM pipeline were both analysed using our DiLiPop tool. Both sets were best fitted by a double-exponential, whose lifetimes are displayed in the inset. (**D**) Comparing the labelling of two tubulin paralogs. (D1) The residence-time distribution (circles) for a typical β-tubulin-labelled embryo is best fitted by a double-exponential similarly to a typical α-tubulin labelled one (Figure 1E). (D2) The lifetimes obtained by analysing (black) $N$ = 25 α-tubulin and (light blue) $N$ = 9 β-tubulin labelled embryos are only mildly different. (**E**) Comparing two different *Caenorhabditis* strains during anaphase. (E1) A double-exponential best fits the residence-time distribution for a typical C. *briggsae* embryo as for C. *elegans* (Figure 1E). (E2) The lifetimes of *C. elegans* ($N$ = 25, black; same data as the ones shown in Figure S2D2) and *C. briggsae* ($N$ = 9, purple) embryos are displayed. In (D) and (E), to detect and link microtubule contacts, we used the KUT pipeline. (**F**) Exemplar NAM analysis of a one-cell embryo. Bright spots are the plus-ends of YFP:: α -tubulin labelled microtubules (left) (Movie S1). They are enhanced after denoising by a ND-SAFIR denoising (middle). The trajectories of the microtubules (yellow lines) are obtained using the ATLAS detection and the MHT linker (right) (Supp. Text, §1.1.3). The parameters used are in Table S4.

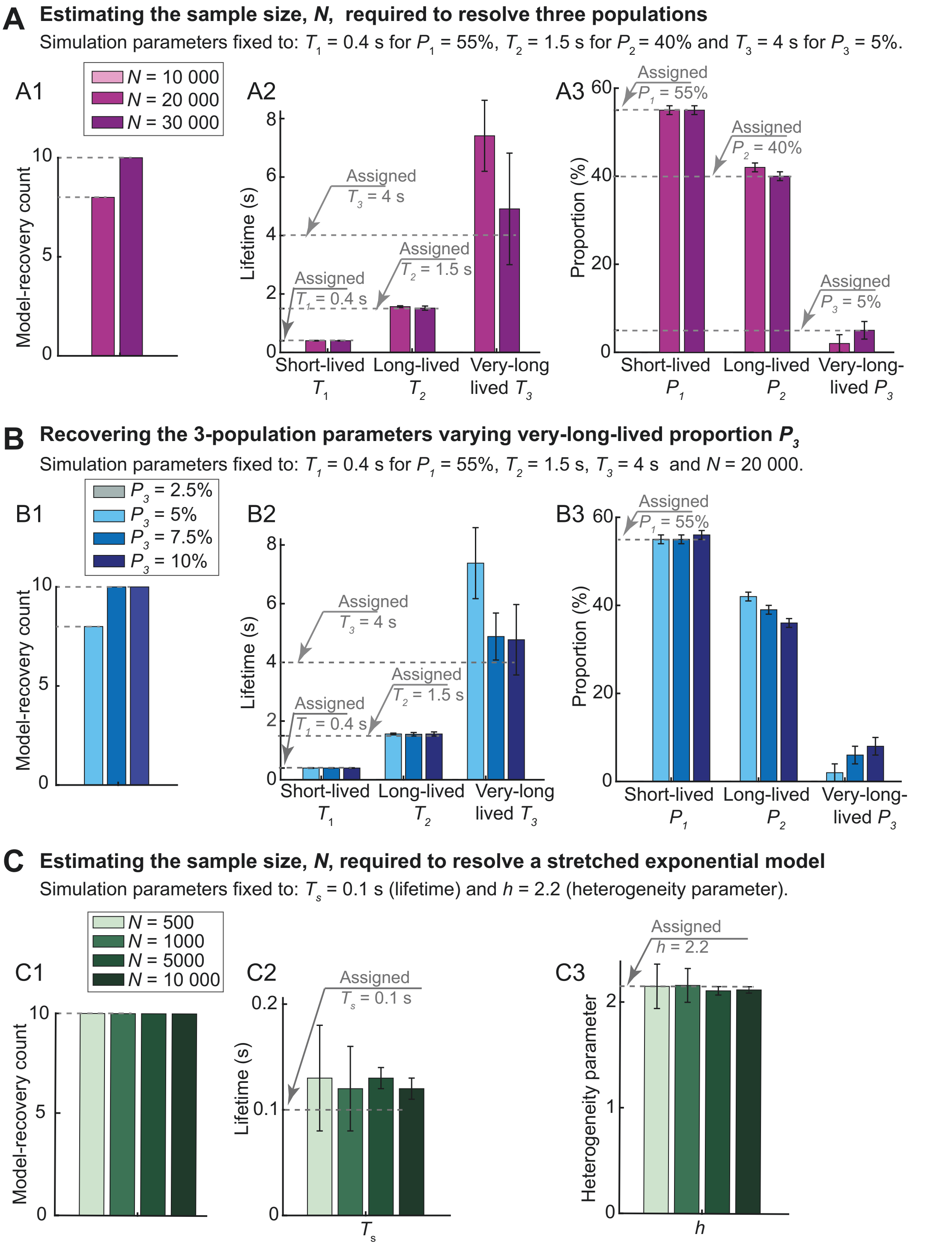


**Supplemental Figure S3: *In silico* investigations reveal the statistical power required to resolve three microtubule populations.**

We analysed *in silico* datasets comprising $Ne$ = 25 "simulated embryos" (see main text) and (**A**) varied the sample size $N$ from 10 000 to 30 000 tracks per embryo while imposing three populations of lifetimes 0.4 s, 1.5 s, 4 s and proportions 55%, 40% and 5%. This process was repeated 10 times, and we report (A1) the number of cases where the triple exponential was the best model. We averaged the (A2) lifetimes and (A3) proportions over these simulations. (**B**) Similar simulations with $N$ = 20 000 tracks per embryo, three populations of lifetimes 0.4 s, 1.5 s and 4 s, with 55% of short-lived tracks, and varying very-long-lived ones between 2.5% and 10%. (**C**) Similarly, we simulated a stretched exponential behaviour, fixing the lifetime to 0.1 s and heterogeneity parameter to 2.2. We varied the sample size *N* between 500 and 10 000 tracks per embryo.

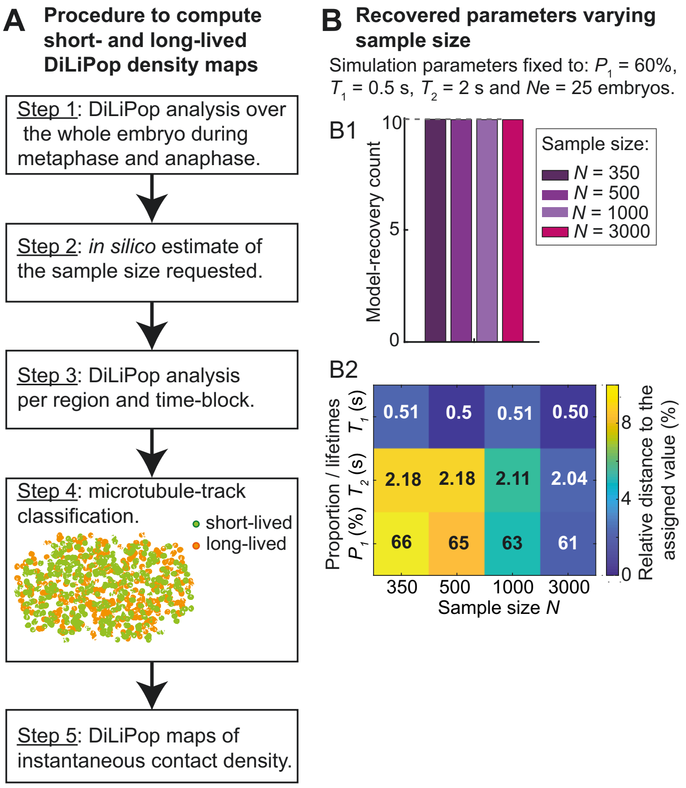


**Supplemental Figure S4: Generating the DiLiPop density maps enables to depict population variations in time and space.**

(**A**) Mapping the short- or long-lived instantaneous contacts and computing the density maps per population (Supp. Text §1.3). (Step 1) We performed a DiLiPop analysis over the whole embryo during metaphase and anaphase. We obtained estimates of the best model and its parameters. (Step 2) From these lifetimes and proportions, we estimated *in silico* the minimal sample size required (see (B) for example). (Step 3) On this ground, we set the size of the regions along the anteroposterior (AP) axis and the time-blocks. We separately DiLiPop-analysed each set of blocks/regions (see Figure 4A1-2, for example). (Step 4) For each microtubule contact, we estimated its probability of belonging to a population, using a single exponential distribution with corresponding-population parameters (as in Figure 1E, Supp. Text §1.2.2). (Step 5) Finally, we computed the instantaneous contact density of each population, using 10 regions along the AP axis and a 10-s running time-window. (**B**) Exemplar estimating of the sample size required to resolve two populations with $Ne$ = 25 simulated embryos, given two dynamical behaviours with lifetimes $T_1$ = 0.5 s and $T_2$ = 2 s and proportions $P_1$ = 60% and $P_2$ = 40%, respectively. We varied the sample size per embryo from 350 to 3000 and repeated 10 times the simulation procedure for each condition. (B1) Number of simulations (out of 10) that recovered the double-exponential as the best model. (B2) Averages of the model parameters recovered over the simulations finding the double-exponential as the best model. The colour shading sets how close the recovered value is from the set one.

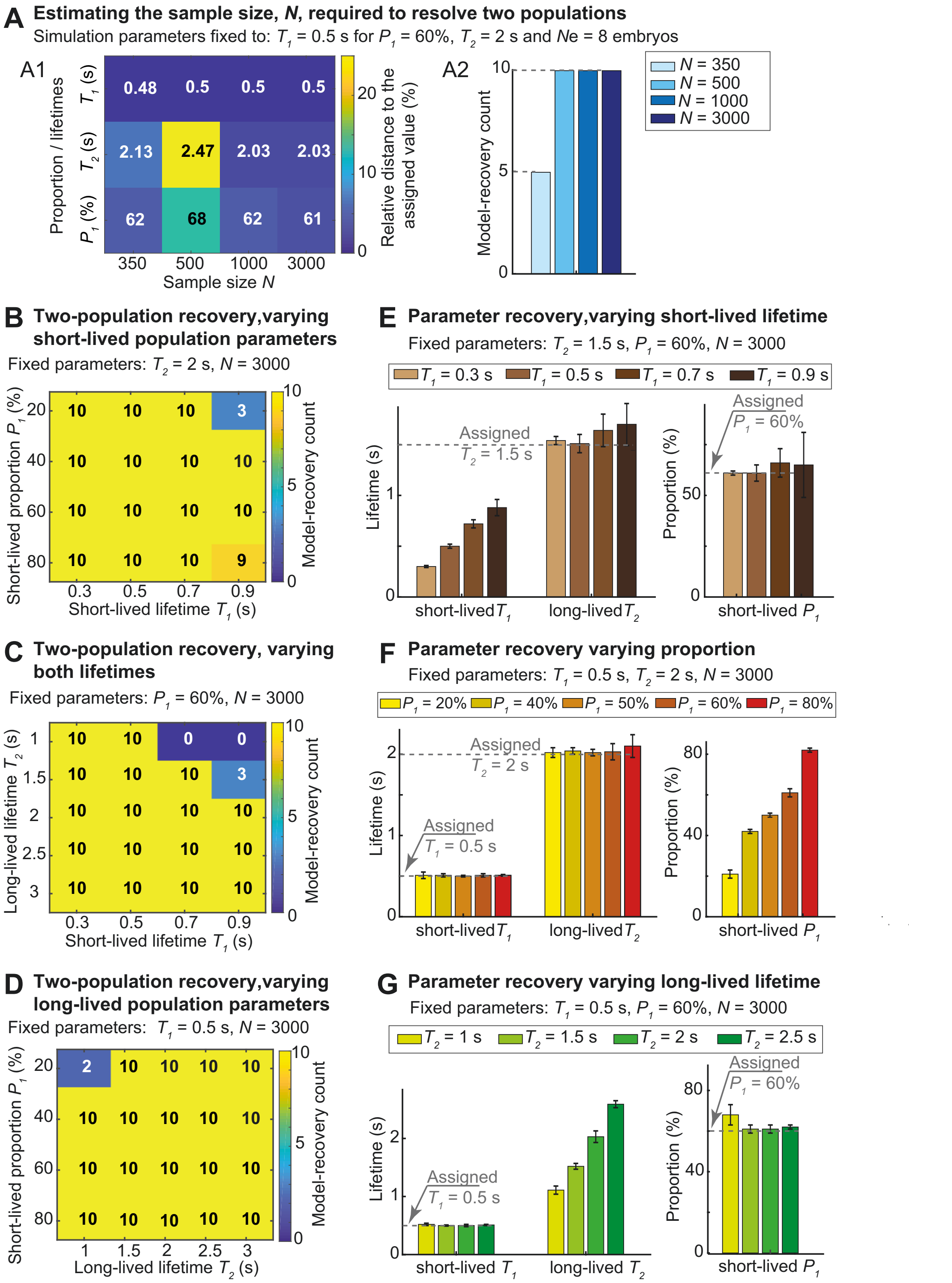


**Supplemental Figure S5: *In silico* investigations estimate the requirements for the DiLiPop-statistical analysis to resolve two populations.**

We analysed *in silico* datasets comprising *Ne* = 8 "simulated embryos" (see main text) and (**A**) varied the sample size *N* from 350 to 3000 tracks per embryo, while imposing two populations of lifetimes 0.5 s and 2 s, with proportions of 60% and 40%. This process was repeated 10 times, and we report (A1) the number of cases where the double-exponential was the best model. We averaged the (A2) lifetimes and (A3) proportions over these simulations. (**B-G**) Similar simulations with *N* = 3000 tracks per embryo, varying: (B) the short-lived lifetime $T_1$ and proportion $P_1$ keeping long-lived lifetime $T_2$ = 2 s; (C) the both lifetimes $T_1$ and $T_2$ keeping short-lived proportion $P_1$ = 60%; (D) the long-lived lifetime $T_2$ and proportion $P_2$ = 1-$P_1$ keeping short-lived lifetime $T_1$ = 0.5 s; (E) the short-lived lifetime $T_1$ keeping the long-lived population parameters $T_2$ = 1.5 s and 1-$P_1$ = 40%; (F) the short-lived proportion $P_1$ keeping the lifetimes $T_1$ = 0.5 s and $T_2$ = 2 s; (G) the long-lived lifetime $T_2$ keeping the short-lived population parameters $T_1$ = 0.5 s and $P_1$ = 60% (Supp. Text, §2).

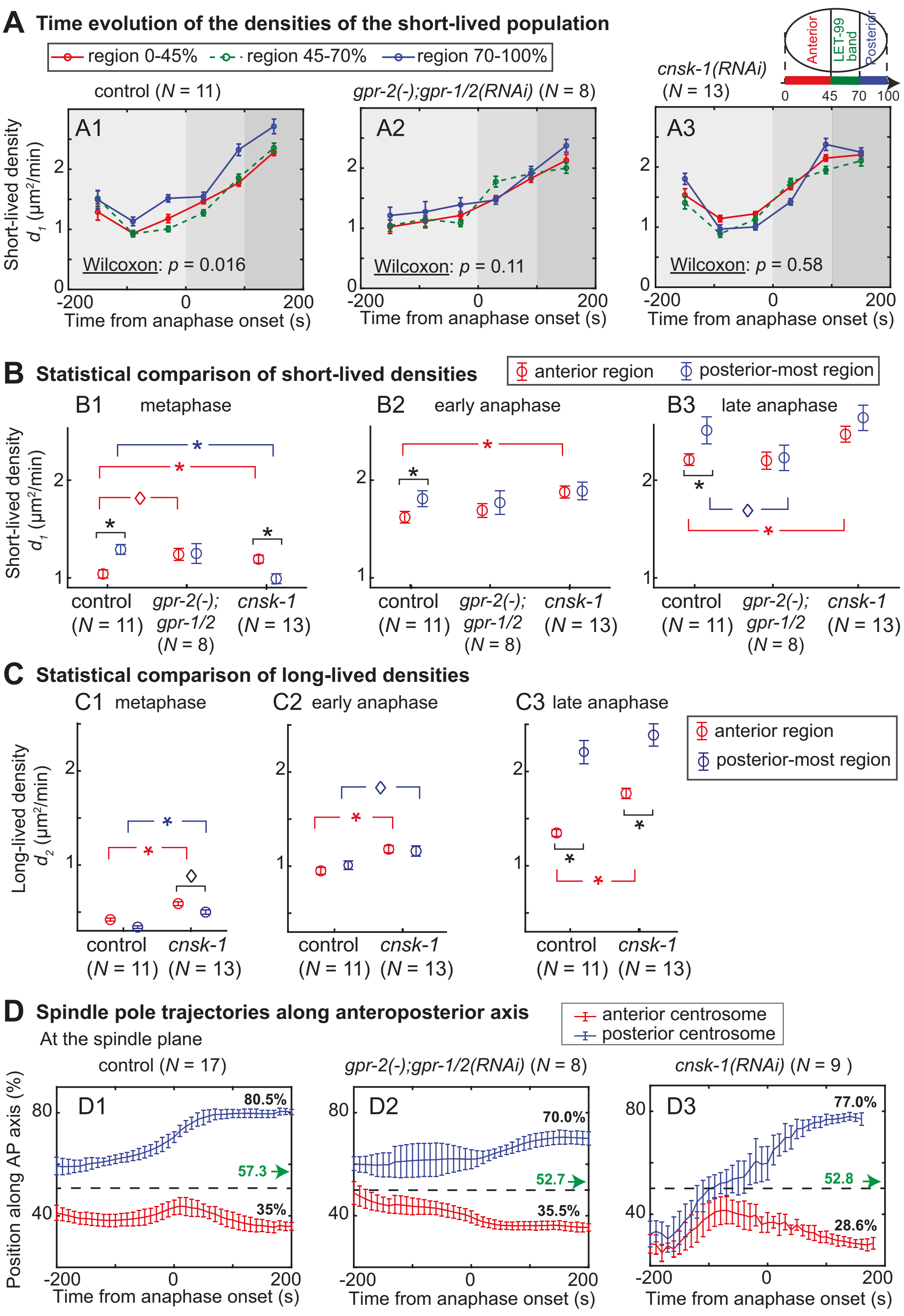


**Supplemental Figure S6: The asymmetry of the force-generator density correlates with the pulling-force-generator distribution.**

(**A**) Evolution of the short-lived densities during metaphase and anaphase in (red) the anterior region, (green) the lateral LET-99 band and (blue) the posterior-most region. These regions are depicted in the schematics at the top right. We analysed, using 60-s time-blocks, (A1) $N = 11$ control embryos, (A2) $N = 8$ *gpr-2(ok1179);gpr-1/2(RNAi)*-treated embryos and (A3) $N = 13$ *cnsk-1(RNAi)*-treated embryos. Standard deviations were computed by the bootstrapping (Supp. Text, §1.2.5). We found a significant difference between the anterior and posterior-most time-series only for the control embryos using the Wilcoxon signed-rank. The grey shadings depict, from lighter to darker, the three time-periods: metaphase (the 200 s before anaphase onset); early anaphase (the 100 s after anaphase onset); and late anaphase (from 100 s to 200 s after anaphase onset). (**B**) The same quantities were analysed, reducing the time resolution to the above three time-periods for the sake of result accuracy. (**C**) Long-lived densities over the three time-periods upon *cnsk-1(RNAi)* (same embryos as in (A-B)). Stars or diamond indicate significant differences (Student's *t*-test). (**D**) The same strains imaged in the same conditions at the spindle plane showed (D2) a reduced posterior displacement for $N$ = 8 *gpr-2(ok1179);gpr-1/2(RNAi)*-treated embryos (6.8 ± 2.1 µm, $p$ = $1.2\text{x}10^{-4}$) and (D3) an increased posterior displacement for $N$ = 11 *cnsk-1(RNAi)*-treated embryos (14.4 ± 4.0 µm, $p$ = $9.2\text{x}10^{-3}$), compared to (D1) $N$ = 17 control embryos (11.7 ± 2.5 µm). The green arrows indicate the final position of the mitotic spindle, while the values in black colour report the final position of the centrosomes.

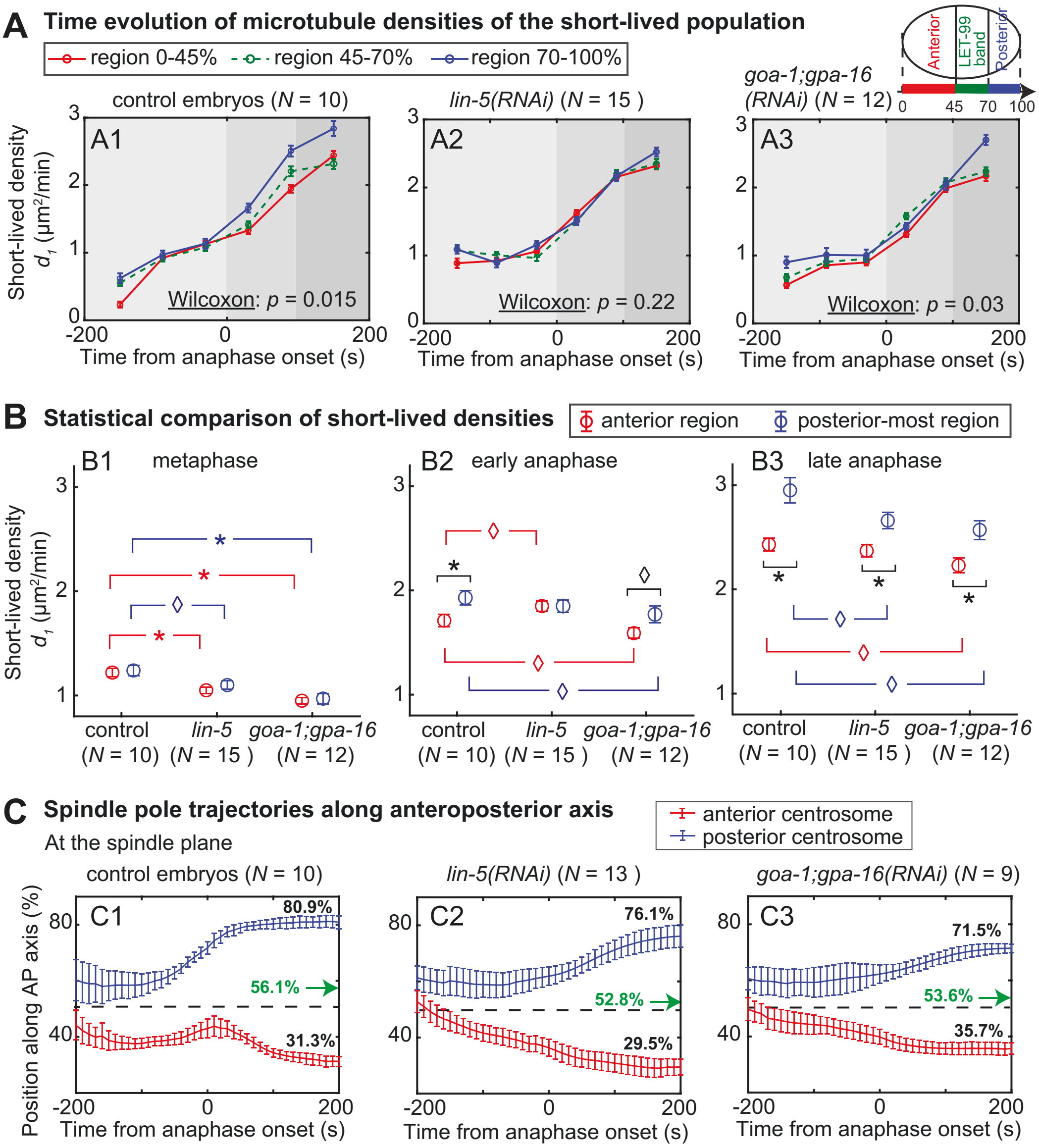


**Supplemental Figure S7: The asymmetry of the short-lived microtubule density reflects the posterior enrichment of the trimeric force-generating complex.**

(**A**) Evolution of the short-lived densities during metaphase and anaphase in (red) the anterior region, (green) the lateral LET-99 band and (blue) the posterior-most region. These regions are depicted in the schematics at the top right. We analysed, using 60-s time-blocks, (A1) $N$ = 10 control embryos, (A2) $N$ = 15 *lin-5(RNAi)*-treated embryos and (A3) $N$ = 12 *goa-1;gpa-16(RNAi)*-treated embryos. Standard deviations were computed by the bootstrapping (Supp. Text, §1.2.5). We found a significant difference between the anterior and posterior-most time-series in control and mildly upon *goa-1;gpa-16(RNAi)* (Wilcoxon signed-rank test). The grey shadings depict, from lighter to darker, the three time-periods: metaphase (the 200 s before anaphase onset); early anaphase (the 100 s after anaphase onset); and late anaphase (from 100 s to 200 s after anaphase onset). (**B**) The same quantities were analysed reducing the time resolution to the above three time-periods for the sake of result accuracy. Star or diamond indicates significant differences (Student's $t$-test) (**C**) The same strain imaged in the same conditions at the spindle plane shows a reduced posterior displacement for (C2) $N$ = 13 *lin-5(RNAi)*-treated embryos (8.1 ± 2.1 µm, $p$ = 1.7x10$^{-3}$) and (C3) $N$ = 9 *goa-1;gpa-16(RNAi)*-treated embryos (7.3 ± 2.6 µm, $p$ = 7x10$^{-4}$), compared to (C1) $N$ = 10 control embryos (11.7 ± 2.5 µm). The green arrows indicate the final position of the mitotic spindle, while the values in black report the final position of the centrosomes.

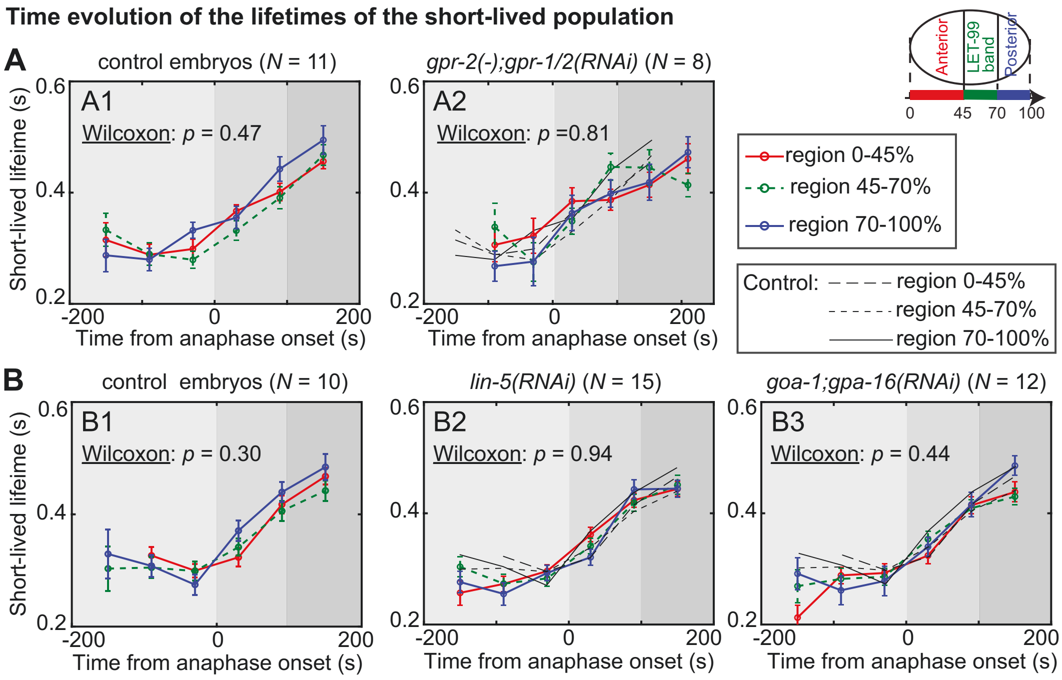


**Supplemental Figure S8: Depleting members of the trimeric force-generating complex does not affect the time evolution of the short-lived microtubule lifetime.**

(**A-B**) Evolution of the short-lived lifetimes during metaphase and anaphase in (red) the anterior region, (green) the lateral LET-99 band and (blue) the posterior-most region. These regions are depicted in the schematics at the top right. We analysed, using 60-s time-blocks, **(A)** on the one hand (A1) $N$ = 11 control embryos, and (A2) $N$ = 8 *gpr-2(ok1179);gpr-1/2(RNAi)*-treated embryos and **(B)** on the other hand (B1) $N$ = 10 control embryos, (B2) $N$ = 15 *lin-5(RNAi)*-treated and (B3) $N$ = 12 *goa-1;gpa-16(RNAi)*-treated embryos. Standard deviations were computed by the bootstrapping (Supp. Text, §1.2.5). We found a significant difference between the anterior and posterior-most time-series in no condition using the Wilcoxon signed-rank test. In (A2), (B2) and (B3), thin black lines report the corresponding controls. The grey shadings depict, from lighter to darker, the three time-periods: metaphase (the 200 s before anaphase onset); early anaphase (the 100 s after anaphase onset); and late anaphase (from 100 s to 200 s after anaphase onset).

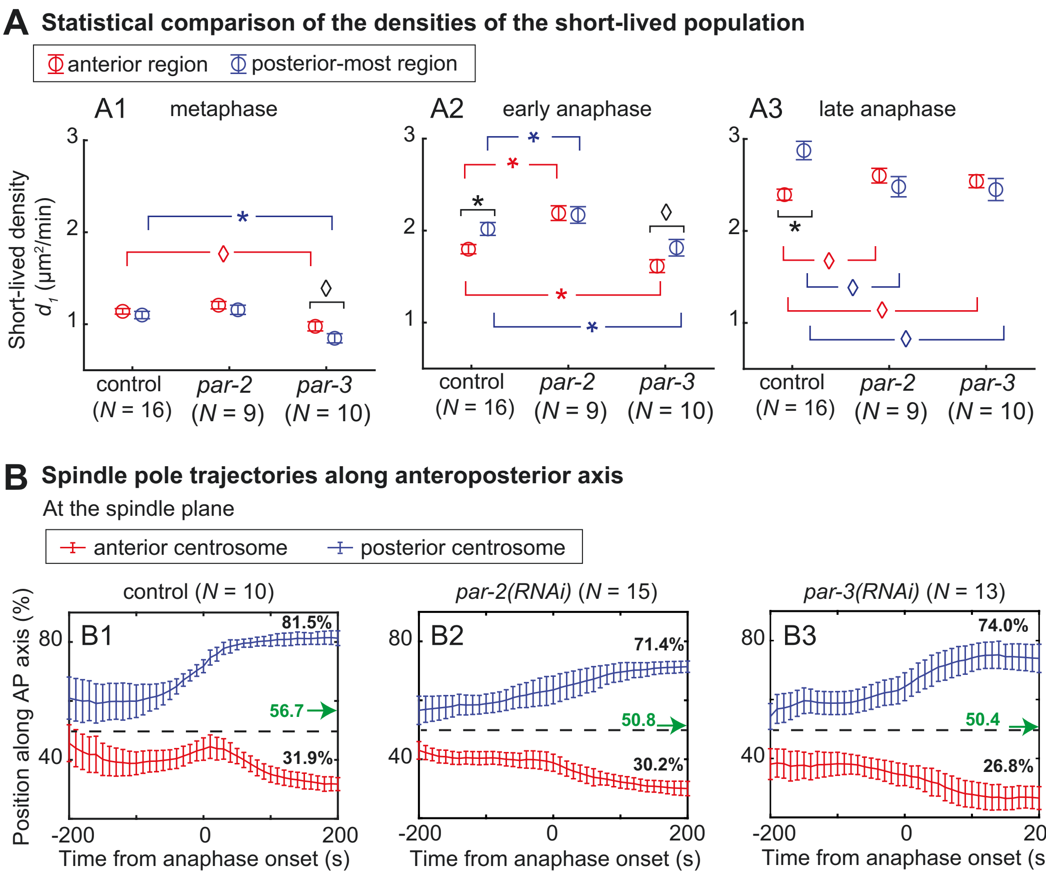


**Supplemental Figure S9: The asymmetry of force-generator density depends on PAR polarity proteins.**

(**A**) Short-lived densities within the two extreme regions for $N$ = 16 control embryos, $N$ = 9 *par-2(RNAi)*-treated embryos and $N$ = 8 *par-3(RNAi)*-treated embryos. We used the same data as in Figure 5A but reduced the time resolution to three time-periods for the sake of result accuracy: (A1) metaphase (the 200 s before anaphase onset); (A2) early anaphase (the 100 s after anaphase onset); and (A3) late anaphase (from 100 s to 200 s after anaphase onset). Stars or diamond indicate significant differences (Student’s *t*-test). (**B**) The same strains imaged in the same conditions at the spindle plane showed a reduced posterior displacement for (B2) $N$ = 15 *par-2(RNAi)*-treated embryos (7.9 ± 1.8 µm, $p = 7\text{x}10^{-4}$) and (B3) $N$ = 13 *par-3(RNAi)*-treated embryos (8.9 ± 1.8 µm, $p = 4\text{x}10^{-3}$), compared to (B1) $N$ = 10 control embryos (12.6 ± 2.5 µm). The green arrows indicate the final position of the mitotic spindle while the values in black colour report the final position of the centrosomes.

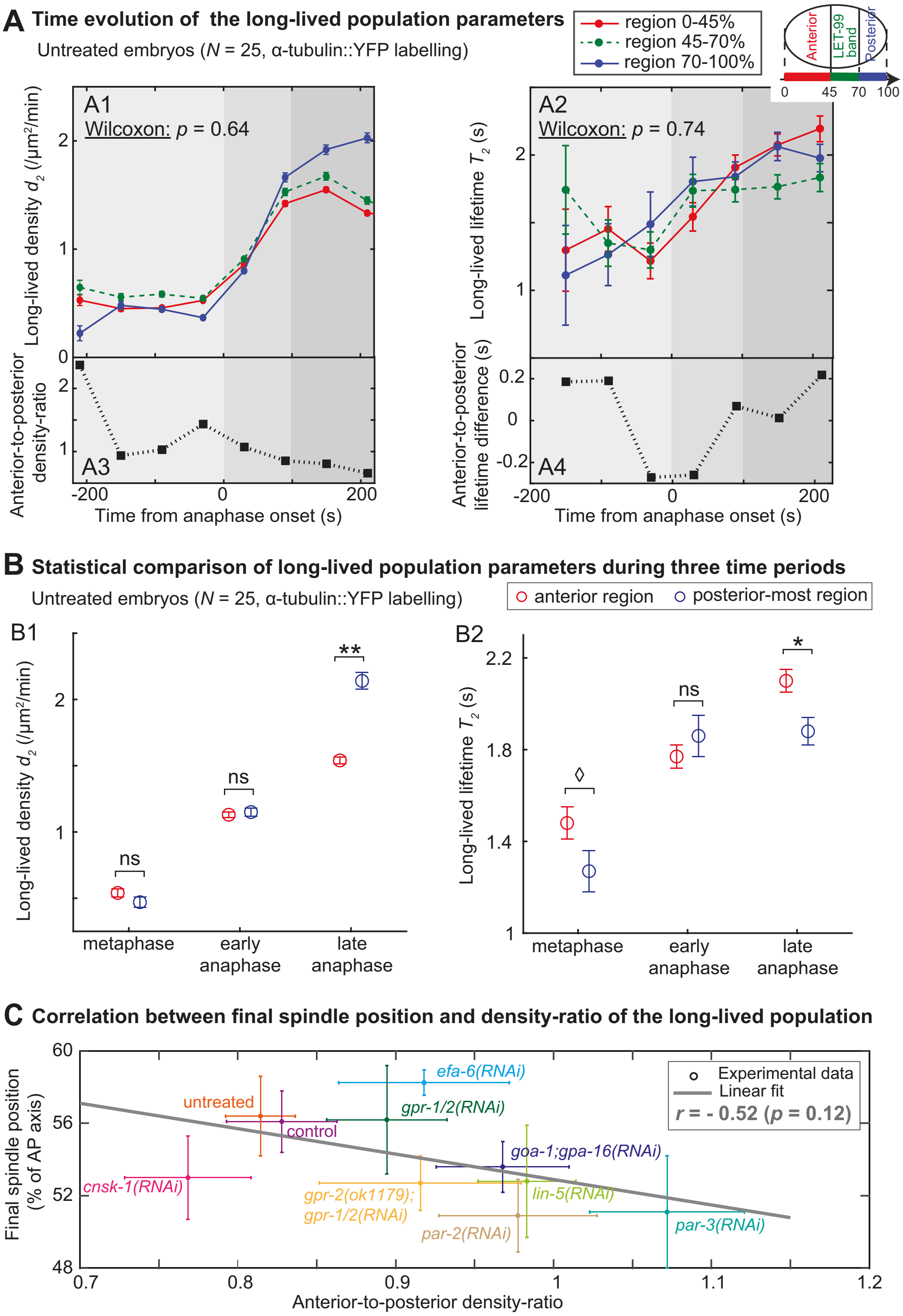


**Figure S10: The long-lived microtubule densities are symmetric until mid-anaphase.**
(**A**) Evolution of the long-lived population (A1) density and (A2) lifetime, during metaphase and anaphase, in (red) the anterior region, (green) the lateral LET-99 band and (blue) the posterior-most region. These regions are depicted in the schematics at the top right. They correspond to the same data as in Figures 2A, 4AB and 6A, i.e. $N = 25$ untreated α-tubulin-labelled embryos but using 60-s time-blocks. Below each plot, either (A3) the anterior-to-posterior density-ratio or (A4) the anterior-to-posterior lifetime differences are plotted. Standard deviations were

computed by the bootstrapping approach (Supp. Text, §1.2.5). We found no significant difference between the time-series (Wilcoxon signed-rank test). The grey shadings depict, from lighter to darker, the three time-periods: metaphase (the 210 s before anaphase onset); early anaphase (the 100 s after anaphase onset); and late anaphase (from 100 s to 210 s after anaphase onset). (**B**) We repeated the same analysis reducing the time resolution for the sake of result accuracy. Stars or diamond indicate significant differences (Student’s *t*-test). (**C**) Final spindle position obtained by imaging the same strain at the spindle plane, plotted against the anterior-to-posterior density ratio for the long-lived population, assessed during the whole anaphase. The grey line depicts the best linear fit, suggesting no correlation. The density ratio was varied by depleting various proteins: *par-3(RNAi)* ($N$ = 10 embryos acquired at the cortex and $N$ = 13 at the spindle plane, further written 10/13), *par-2(RNAi)* ($N$ = 9/16), *gpr-2(ok1179);gpr-1/2(RNAi)* ($N$ = 8/8), *cnsk-1(RNAi)* ($N$ = 13/9), *lin-5(RNAi)* ($N$ = 13/14), *goa-1;gpa16(RNAi)* ($N$ = 12/9), *gpr-1/2(RNAi)* ($N$ = 11/6), *efa-6(RNAi)* ($N$ = 10/11), $N$ = 11/10 control embryos, and $N$ = 25/9 untreated embryos. Error bars are the standard deviations.

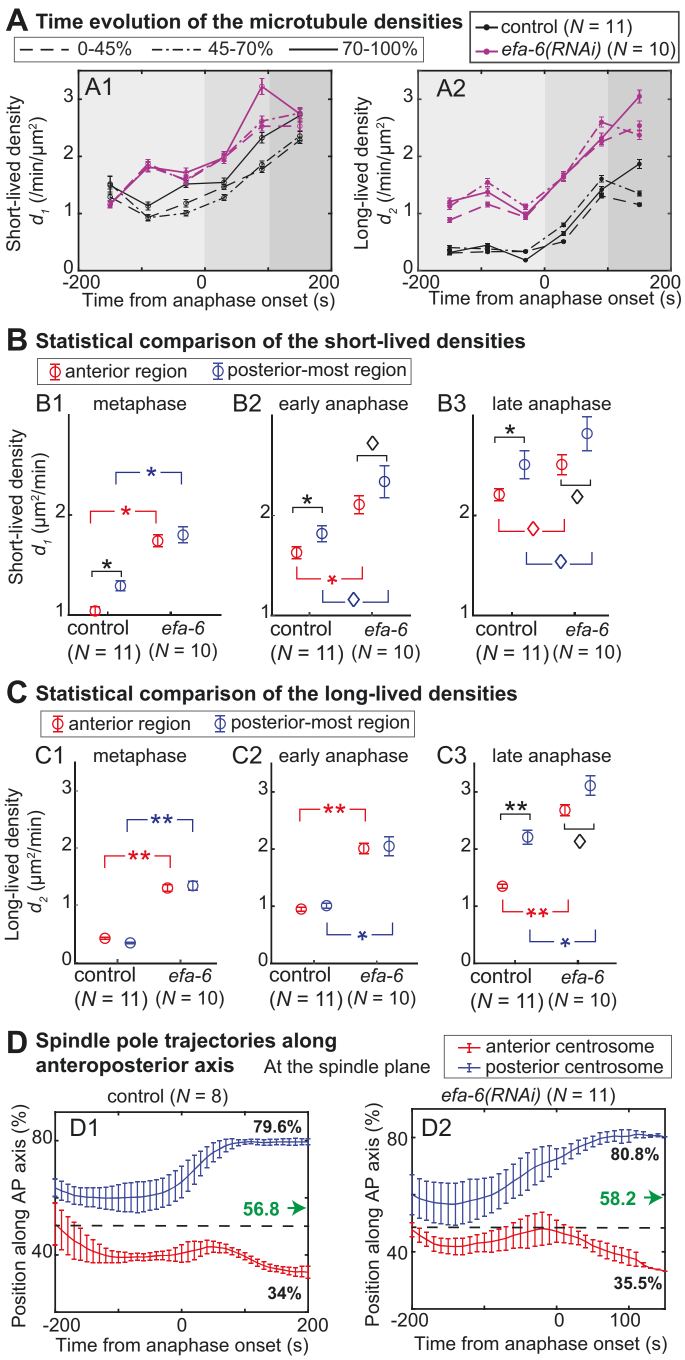


**Supplemental Figure S11: The pushing related microtubule population does not contribute to the spindle posterior-displacement.**

(**A**) Evolution of the (A1) short-lived and (A2) long-lived densities during metaphase and anaphase, using 60-s time-blocks, in the three cortical regions for (black) $N$ = 11 control embryos and (purple) $N$ = 10 *efa-6(RNAi)*-treated embryos. Standard deviations were computed by the bootstrapping (Supp. Text, §1.2.5). The grey shadings depict, from lighter to darker, the three time-periods: metaphase (the 200 s before anaphase onset); early anaphase (the 100 s after anaphase onset); and late anaphase (from 100 s to 200 s after anaphase onset). (**B-C**) We repeated the same analysis reducing the time resolution for the sake of result accuracy. Stars or diamond indicate significant differences (Student's $t$-test). (**D**) The same strains imaged in the same conditions at the spindle plane showed a similar posterior displacement for (D2) $N$ = 11 *efa-6(RNAi)*-treated embryos (13.6 ± 5.3 µm, $p$ = 0.27) compared to (D1) $N$ = 8 control embryos

(11.3 ± 3.2 μm). The green arrows indicate the final position of the mitotic spindle while the values in black colour report the final position of the centrosomes.

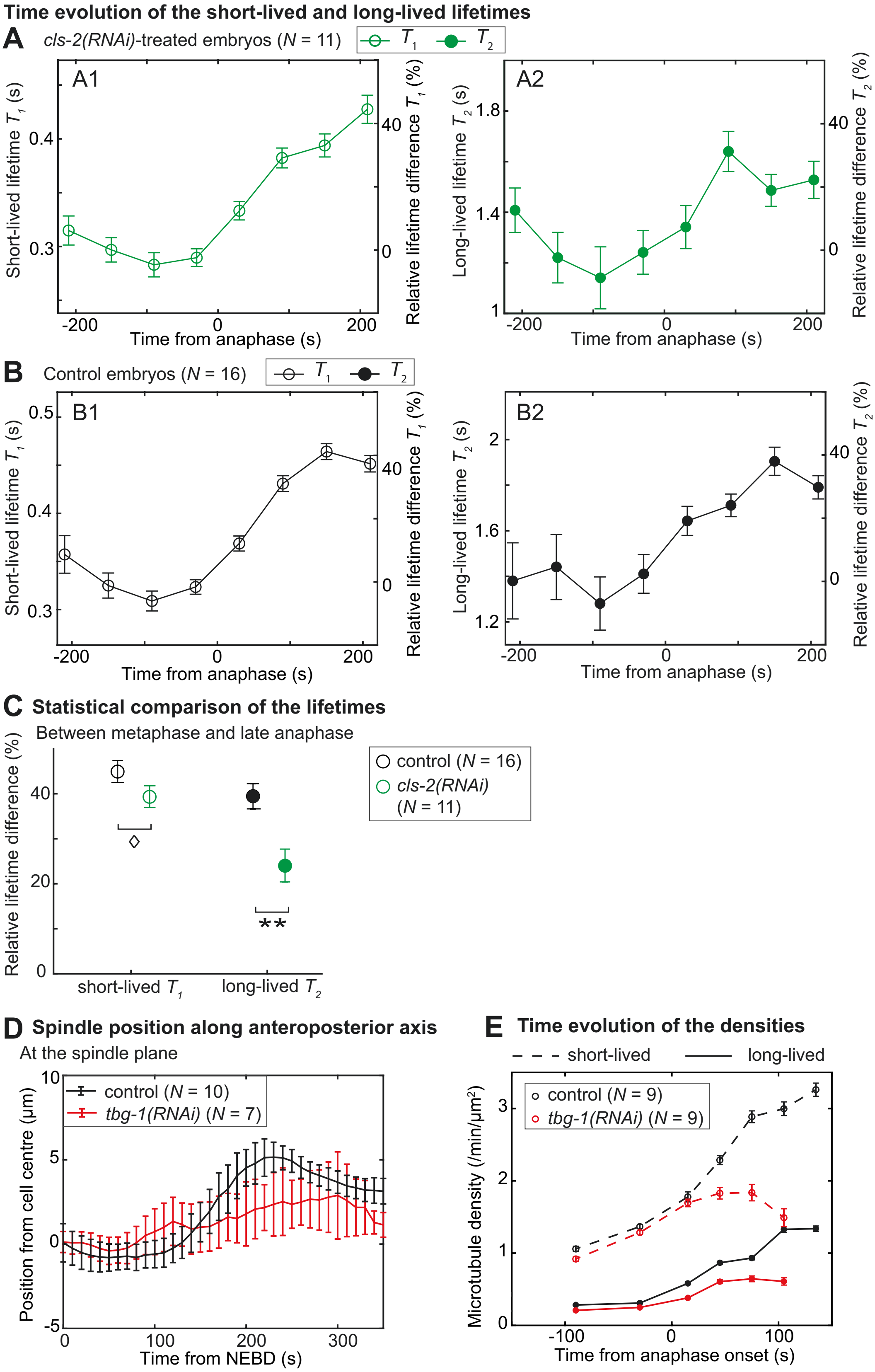


**Supplemental Figure S12: The long-lived microtubule lifetime depends on CLS-2$^{CLASP}$.**
(**A-B**) Temporal evolutions of the microtubule lifetimes of (**A**) (green) $N$ = 11 *cls-2(RNAi)*-treated embryos for (A1) short-lived and (A2) long-lived populations, and (**B**) (black) their control embryos ($N$ = 16). We considered a single region encompassing the whole cortex and

used 60-s time-blocks during metaphase and anaphase. Right Y-scale displays relative lifetime difference from metaphase mean value. The long-lived-lifetime time-series of the control and *cls-2(RNAi)*-treated embryos were mildly independent (Pearson $r$ = 0.78, $\chi^2$ test $p$ = 0.022), while the short-lived-lifetime ones were correlated ($r$ = 0.97, $p$ = 4.6x$10^{-5}$). (**C**) Relative differences in short-lived and long-lived lifetimes between metaphase and late anaphase (100 s to 200 s from anaphase onset), normalised by their respective metaphase lifetimes, for (green) $N$ = 11 *cls-2(RNAi)*-treated embryos and (black) $N$ = 16 control embryos. Standard deviations were obtained by bootstrapping in plots (A-C) (Supp. Text, §1.2.5). Stars or diamond indicate significant differences (Student’s $t$-test). (**D**) Position of the mitotic spindle during mitosis for (red) $N$ = 7 *tbg-1(RNAi)*-treated embryos and (black) $N$ = 10 control embryos. (**E**) Temporal evolution of the microtubule densities of (dashed line) the short-lived and (plain line) long-lived populations in (red) $N$ = 9 *tbg-1(RNAi)*-treated embryos and (black) $N$ = 9 control embryos. We considered a single region encompassing the whole cortex and used 60-s time-blocks during metaphase and 30-s time-blocks during anaphase. Error bars were obtained by bootstrapping (Supp. Text, §1.2.5).

| | |
|---|---|
| **Denoising (Kalman filter)** | |
| Gain | 0.5 |
| Initial estimate of the noise | 0.05 |
| **Detection (u-track algorithm)** | |
| Gaussian standard deviation | 1.2 |
| Alpha-value for initial detection of local maxima | 0.14 |
| Rolling window time-averaging | 3 |
| Iterative Gaussian mixture-model fitting | 0 |
| **Tracking (u-track algorithm)** | |
| Maximum gap to close | 5 |
| Merge split | 0 |
| Minimum length of track segments from the first step | 1 |
| **Cost function frame-to-frame linking (u-track algorithm)** | |
| Flag for linear motion | 1 |
| Allow instantaneous direction reversal | 1 |
| Search radius lower limit | 1 |
| Search radius upper limit | 3 |
| Standard deviation multiplication factor | 3 |
| Nearest neighbour distance calculation | 1 |
| Number of frames for nearest neighbour distance calculation | 4 |
| **Cost function close gaps (u-track algorithm)** | |
| Flag for linear motion | 1 |
| Search radius lower limit | 1 |
| Search radius upper limit | 3 |
| Standard deviation multiplication factor | 3 |
| **Nearest neighbour distance calculation (u-track algorithm)** | |
| Number of frames for nearest neighbour | 4 |
| Penalty for increasing gap length | 1.5 |
| Maximum angle between linear tracks segments | 30 |

**Table S1: Parameters used for KUT-pipeline analysis.**
The pipeline KUT, illustrated in Figure 1A, performs sequentially a Kalman denoising and particle tracking through the u-track algorithm.

| **Model** | Mono-exponential | Double-exponential | Triple-exponential | Stretched exponential |
|---|---|---|---|---|
| **BIC value** | 76871 | **19247** | 19353 | 21558 |
| **BIC-derived model probability** | 0 | **1** | 1.3 x $10^{-23}$ | 0 |
| **Fit-estimated lifetimes (s)** | 0.98 | **0.44 / 1.86** | 0.42 / 1.58 / 4.39 | 0.095 |
| **Fit-estimated Proportions (%)** | - | **60.4 / 39.6** | 56.1 / 40.1 / 3.8 | - |
| **Fit-estimated heterogeneity parameter (unit less)** | - | **-** | - | 2.188 |

**Table S2: Typical values of the Bayesian Inference Criterion used to select the best model.**

The distributions of microtubule-track durations for $N$ = 25 untreated *C. elegans* embryo imaged at the cortex (about 20 000 microtubule tracks per embryo) were fitted using four different models (Supp. Text, §1.2.1, same data as Figure 1). The table reproduces the parameters for the four models adjusting the experimental distributions. The minimum value of the Bayesian Inference Criterion (BIC) indicates that (blue) the double-exponential model offers the best fit (see curves and parameters in Figure 1EF).

| | |
|---|---|
| Image size (pixels) | 256 x 256 |
| Duration (frames) | 3000 |
| Density of particles (/µm$^2$) | 1 |
| Quantum yield | 0.61 |
| Pixel size (nm) | 0.139 |
| Sampling rate (s) | 0.1 |
| PSF shape | Gaussian |
| PSF size (µm) | 0.244 |
| Dynamic range of the fabricated images (bits) | 12 |
| Spot diffusion coefficient (µm$^2$/s) | 0.002 |
| Standard deviation of the background noise | 0.5 |
| Mean track duration (s) | 1 (Figure S2A);<br>0.4 and 2 (Figure S2B) |

**Table S3: Parameters used to generate images *in silico*.**
We fabricated images with particles of known dynamics, mimicking the cortex of embryos with entirely fluorescent-labelled microtubules. This *in silico* dataset acts as ground-truth to validate the analysis pipelines.

| | |
|---|---|
| **ND-SAFIR** | |
| Patch size | 3 |
| Iterations | 3 |
| Peak | no |
| **ATLAS** | |
| *p*-value | 0.004 |
| **MHT** | |
| Parameter values | Default ones for diffusive particles |

**Table S4: Parameters used for NAM-pipeline processing.**
The image-processing pipeline NAM, illustrated in Figure S2F, performs sequentially a ND-SAFIR denoising step, an ATLAS spot-detection, and a MHT particle-linking step.

**Movie S1: Fluorescent spots of microtubules contacting the cortex of a *C. elegans* embryo.**

Movie of a *Caenorhabditis elegans* one-cell untreated embryo imaged at the cortex plane. Microtubules were labelled by YFP::α-tubulin and images were acquired at a frequency of 10 Hz. The movie was accelerated to 5x real-time.

**SUPPLEMENTARY TEXT** for the paper entitled “**The coordination of spindle-positioning forces during the asymmetric division of the *C. elegans* zygote is revealed by distinct microtubule dynamics at the cortex.**”

### Table of Contents

**1 The DiLiPop assay faithfully disentangles spots with mixed dynamics. . 2**

**1.1 Fast imaging of microtubule contacts at the cortex and advanced image processing ... 2**

1.1.1 Microscopy modality and imaging conditions ... 2

1.1.2 Tracking of the microtubule cortical contacts in low SNR images using a dedicated image processing ... 2

1.1.3 Validating the two-population characteristics using a different image-processing pipeline. ... 3

**1.2 Revealing the dynamics of the microtubule contacts through advanced statistical analysis. ... 4**

1.2.1 Modelling mixed populations of microtubules with diverse dynamics ... 4

1.2.2 Model fitting using a maximum likelihood estimator based on Poisson-distributed errors ... 5

1.2.3 Global fitting of a set of embryos recorded in the same condition ... 6

1.2.4 Choosing the best fitting model ... 6

1.2.5 Estimating of the confidence interval on the model parameters ... 6

**1.3 Mapping the short- or long-lived instantaneous contacts. ... 7**

**2 Establishing the requirements to distinguish various microtubule populations. ... 9**

### 1 The DiLiPop assay faithfully disentangles spots with mixed dynamics.

We developed a method to measure the dynamics of the microtubules at the cortex. We applied it here to understand the regulation of the spindle-positioning forces in space (along anteroposterior (AP) axis) and time (throughout mitosis) in the *C. elegans* one-cell embryo. We could distinguish various populations of spots, i.e. microtubule-end contacts, through their distinct dynamical behaviours. The sensibility is good enough to enable a spatiotemporal resolution appropriate to study the regulation of each of these populations. Our so-called DiLiPop ("*Distinct Lifetime sub-Population*") assay is technically detailed in this Supplementary Text. The workflow encompasses five steps: (a) to image the microtubule contacts at the cortex at a high frame rate (10 frames/s); (b) to track these contacts by advanced image processing (Figure 1A); (c) to classify the tracks according to their duration (Figure 1B); (d) to fit the duration distribution using finite-mixture-of-exponential models by global maximum likelihood (Figure 1C,E); and finally (e) to select the model that best fits the experimental distribution using the Bayesian inference. The figure 1D schematises this workflow.

#### 1.1 Fast imaging of microtubule contacts at the cortex and advanced image processing

##### 1.1.1 Microscopy modality and imaging conditions

To faithfully measure the microtubule-contact dynamics at the cortex throughout mitosis in physiological conditions, we ensured: (a) to preserve the embryo shape as much as possible since it influences the spatial distribution of microtubule contacts at the cortex (Bouvrais *et al.*, 2018); (b) to acquire at a high enough rate to correctly capture the fast microtubule dynamics (Srayko *et al.*, 2005; Kozlowski *et al.*, 2007; O'Rourke *et al.*, 2010; Schmidt *et al.*, 2017; Bouvrais *et al.*, 2018; Sugioka *et al.*, 2018); and (c) to control and limit the photo-bleaching and photo-damaging during the typical 5 minutes of imaging as nematode zygotes are known to be sensitive (Tinevez *et al.*, 2012). Practically, we imaged at the cortical plane, using an optimised spinning disk microscope to cope with the thickness of the perivitelline space of about 250-750 nm (Olson *et al.*, 2012). We used an EMCCD camera, whose sensitivity enabled us to image at 10 frames/s – fast enough to resolve dynamics and with exposure long enough to ensure a signal-to-noise ratio (SNR) appropriate for our image-processing pipelines.

##### 1.1.2 Tracking of the microtubule cortical contacts in low SNR images using a dedicated image processing

We tracked the microtubule-contact spots to determine their residence times at the cortex. Compared to bare spot detection on single frames, it provides robustness when used on limited-contrast images (Bouvrais *et al.*, 2018). The low signal-to-noise ratio combined with fast dynamics makes the tracking a tour-de-force. It requires tailored processing, especially to distinguish populations of close but distinct dynamics (Coelho *et al.*, 2013). Our analysis pipeline follows the classic three steps: to denoise the images applying Kalman filter (Kalman, 1960) (Figure 1A, middle); to detect the spots using u-track (Jaqaman *et al.*, 2008); and to link the candidates to produce tracks of microtubule contacts using once again u-track (Figure 1A, bottom). This analysis pipeline is further named *KUT pipeline* and parameters used are found in Table S1. To fully interpret these tracks, we recovered the shape of the embryo adhesion on the cover-slip taking advantage of the cytoplasmic fraction of the labelled tubulin and using active contours (Pécréaux *et al.*, 2006). It enabled us: (a) to define a temporal reference set as the cytokinesis-furrow ingression onset; (b) to get the area of the embryo across mitosis to measure the density of microtubule contacts; and (c) to select real tracks, lying within the embryo, by masking.

We obtained tens of thousands of microtubule tracks for each embryo distributed from the NEBD to late anaphase (Figure 1B). As a consistency check, we computed, for each embryo, the number of instantaneous contacts, i.e. present in any single frame, and observed its time variation (Figure S1A). We found an excellent reproducibility between embryos. We also recovered an increase at anaphase onset, expected from the change in microtubule dynamics (Srayko *et al.*, 2005). This increase continued until late anaphase, likely because the centrosome-to-cortex distance is decreasing (Bouvrais *et al.*, 2018). Extrapolating this count to the whole cortex, we estimated that our imaging and analysis pipeline captured at least 2/3 of the expected total number of microtubule contacts based on electron tomography (Redemann *et al.*, 2017). Our sampling is, therefore, very representative of the whole population.

##### 1.1.3 Validating the two-population characteristics using a different image-processing pipeline.

While the choice of these algorithms appears sensible, in particular, to link dense spots into short tracks, we set to ensure that the biological results are not dependent on this choice of tools. Furthermore, we reckoned that the microtubule lifetimes, i.e. the duration of the tracks, are related to the ability of the image processing to detect weak spots and link them between frames continuously. Therefore, we built a second image-processing pipeline, further named *NAM*. In this pipeline, ND-SAFIR algorithm denoises the images (Boulanger *et al.*, 2010), ATLAS ("adaptive thresholding of Laplacian of Gaussian images with auto-selected scale") ensures spot detection (Basset *et al.*, 2015), and MHT (Multiple Hypothesis Tracker) preforms the linking (Chenouard *et al.*, 2013) (Figure S2F). The parameters used are reproduced in Table S4. Processing *in silico* images mimicking microtubule contacts at the cortex enabled to optimise those parameters (Material and Methods) (Costantino *et al.,* 2005). Importantly, each step relies on different approaches compared to the ones composing the KUT pipeline. Indeed, the denoising by the Kalman filter in the KUT pipeline acts recursively, using temporal information to discriminate signal from noise, while ND-SAFIR is a patch-based adaptive method, using both space and time. The detection by u-track requires the adjustment of several parameters (e.g. background level, Point Spread Function (PSF), frame averaging), while ATLAS requires only a threshold *p*-value. Finally, the principle of u-track linker is a linear assignment problem which optimizes a cost matrix globally in space and mostly locally in time, while the other linker uses an iterative multi-frame tracking procedure with a maximisation of the track likelihood. We repeated the analysis of $N = 20$ untreated embryos. For each embryo, the track-duration distributions appeared very similar to the KUT pipeline one (Figure S2C). Over the whole set, we also found two dynamically distinct microtubule populations and their lifetimes are close to KUT-pipeline ones (Figure S2C, inset). Interestingly, the differences affect mostly the long tracks (Figure S2C), and they are likely related to the ability to track faint dots (the longer they last, the more they are photo-bleached) and to link them. However, the very close results, using two different image-processing pipelines reinforced the presence of two populations of microtubules contacting the cortex, distinct by their dynamics.

#### 1.2 Revealing the dynamics of the microtubule contacts through advanced statistical analysis.

To investigate the dynamical behaviours of the microtubule tracks at the cortex, we classified them based on the histogram of their residence times (Figure 1B). In particular, we set to disentangle mixed dynamics reflected by different residence times. Such an aim was challenging because the various populations exhibit close times, and the track durations were short. We developed a methodology in three steps (Figure 1D): (a) to estimate the model parameters (Figure 1D, orange shading), (b) to select the best model among mixture-of-exponential ones (Figure 1D, blue shading), and (c) to estimate the errors associated to the model parameters (Figure 1D, purple shading).

##### 1.2.1 Modelling mixed populations of microtubules with diverse dynamics

The dynamics of the microtubules at the cortex are recapitulated by the rate of growing microtubule switching to depolymerisation state, termed catastrophe (Cassimeris *et al.*, 1988). It is the inverse of the cortical residence time or lifetime used here. Catastrophe is a stochastic process, appropriately modelled as an exponential decay (Fygenson *et al.*, 1994). *In vivo*, microtubule dynamics are altered by numerous microtubule-associated proteins (MAP). In particular, they can stabilise the microtubules at the cortex, decreasing their catastrophe rate, or foster depolymerising. Consequently, different catastrophe rates may co-exist in a population of microtubules, leading to a residence time distribution well approached by a mixture-of-exponential model. We tested three distinct models: mono-exponential (Figure 1C, blue line), double-exponential (Figure 1C, red line) or triple-exponential (Figure 1C, green line) corresponding respectively to a single population, or the co-existence of two or three microtubule populations. The *in silico* tests suggested that it is not possible to distinguish more populations in our experimental conditions, given the number of contacts detectable and the accessible frame rates (main text, p. 7; Figures S3A, S4B). We also considered the case where the microtubules display a broad range of catastrophe rates without distinguishable population. We modelled this case by a stretched exponential (Figure 1C, purple line). The equations corresponding to these different models are reproduced below:

**Mono-exponential model**:

$$f_{mono} = \frac{M}{R}\,\mathrm{e}^{-t/T} \qquad (1)$$

with $M$ the number of microtubules, $R$ normalises the probability distribution, $t$ the residence time and $T$ the characteristic lifetime of the exponential decay.

**Double-exponential model**:

$$f_{dble} = M\Big[\frac{P_1}{R_1}\,\mathrm{e}^{-t/T_1} + \frac{(1-P_1)}{R_2}\,\mathrm{e}^{-t/T_2}\Big] \qquad (2)$$

with $P_1$ the proportion of microtubules in the short-lived population, and $T_1, T_2$ the lifetimes of, respectively, the short-lived and long-lived populations.

**Triple-exponential model**:

$$f_{triple} = M\Big[\frac{P_1}{R_1}\,\mathrm{e}^{-t/T_1} + \frac{P_2}{R_2}\,\mathrm{e}^{-t/T_2} + \frac{(1-P_1-P_2)}{R_3}\,\mathrm{e}^{-t/T_3}\Big] \qquad (3)$$

with $P_1, P_2, P_3 (= 1 - P_1 - P_2)$ the proportions of the short-, mid- and long-lived populations, and $T_1,\ T_2,\ T_3$ the corresponding lifetimes.

**Stretched exponential model**:

$$f_{stretched} = \frac{M}{R_s}\,\mathrm{e}^{-[t/T_s]^{(1/h)}} \qquad (4)$$

with $T_s$ the lifetime of the exponential decay and $h$ the heterogeneity parameter of the sample.

##### 1.2.2 Model fitting using a maximum likelihood estimator based on Poisson-distributed errors

Because we work with a limited number of tracks, the classic implementation of the objective (cost) function using a least-square estimator did not appear sensible. Indeed, it assumes Gaussian-distributed experimental errors (Barford, 1985). In contrast, we reverted to the Maximum-Likelihood Estimator (MLE) and accounted for the discrete nature of the histogram. In such a case, the Poisson distribution is a suitable model of the distribution of the experimental errors, and we used it to design our objective function $L_j$, with $j$ indexing the embryos (Figure 1D, grey shading) (Laurence and Chromy, 2010):

$$-\log(L_j) = \sum_i \hat{c}_l - \sum_i c_{i,j}\log(\hat{c}_l) \qquad (5)$$

$i$ indexes the bins, $c_{i,j}$ the corresponding experimental count of tracks and $\widehat{c_i}$ the model estimate (optimum).

##### 1.2.3 Global fitting of a set of embryos recorded in the same condition

In this paper, we investigate multiple populations of microtubules, distinct by their cortical residence times within a single embryo. We excluded fitting a global distribution obtained cumulating embryos on the one hand. It would leave the opened possibility of a single microtubule population per embryo but with parameters distinct between embryos. On the other hand, the limited number of microtubules per embryo made fitting each embryo separately not accurate enough. We thus used global fitting (Beechem, 1992). It imposes the same model and parameters for each embryo of the data set, but the likelihood is computed independently for each. We maximised the global likelihood combining the individual ones (Figure 1D, green shading):

$$\log(L^{total}) = \sum_{j=1}^{N_e} \log(L_j) \qquad (6)$$

with $Ne$ is the number of embryos. By increasing the amount of data in use, such a strategy increased the precision and reliability of our estimates.

##### 1.2.4 Choosing the best fitting model

To test whether multiple populations of microtubules, distinct by their dynamics, co-exist, we performed a statistical test to decide the best model (Figure 1D, blue shading). To mitigate the risk of overfitting (i.e. improving the fit by adding more free parameters), we used the Bayesian Inference Criterion (*BIC*, Equation 7) (Schwarz, 1978) reading:

$$BIC = \sum_{j=1}^{N_e} -2\log(L_j) + V \times \log(M_j) - 2\log(L_j) \qquad (7)$$

with $V$ the number of free parameters of the model, and $M_j$ the number of microtubules for the embryo $j$.

To gain confidence, we concurrently used the Akaike Information Criterion with a correction for finite sample (*AICc*, Equation 8) (Wagenmakers and Farrell, 2004). In most of the cases (>95%), the two criteria were in agreement. In case of disagreement, we preferred the *BIC,* designed for deciding about statistical significance, in contrast to *AIC* initially developed for predictive purposes.

$$AICc = \sum_{j=1}^{N_e} -2\log(L_j) + 2V + \frac{2V(V+1)}{M_j - V - 1} \qquad (8)$$

##### 1.2.5 Estimating of the confidence interval on the model parameters

To assess the uncertainty on the fit parameters, we used two different methods (Figure

1D, purple shading): (a) the "likelihood ratio", relying on a model of the objective function near the optimum (Figure S1E); (b) the bootstrapping, which offers an empirical estimate, model-free (Figure S1D).

(a) By an approach similar to the **likelihood ratio** test (Bolker, 2008; Agresti, 2013), we compared the optimal likelihood $L_{abs}$ obtained by global fitting and the one $L_{rest}$ obtained when fixing the parameter of interest $p$ and optimizing the others (Figure S1E, brown and red shadings). Varying $p$ enabled us to estimate the likelihood of that parameter around its best-estimate $\hat{p}$. Twice the log of the ratio of the likelihoods $L_{rest}/L_{abs}$ can be modelled by a chi-squared distribution with $r$ degrees of freedom, and $r$ is the number of fixed parameters.
Having two degrees of freedom and setting the confidence level to 95%, the chi-value in our example was equal to 5.99, so that the 95% threshold in $\log(L)$ (purple dashed line) reads:

$$\log(L_{thresh}) = \log(L_{abs}) - 3 \qquad (9)$$

The bounding values of the parameter correspond to the confidence interval (Figure S1E, blue shading). The figure S1BC exemplifies this approach.

(b) The **bootstrapping** approach is based on resampling the experimental data for each embryo (Efron and Tibshirani, 1993). It requires a high amount of data to be precise. Practically, we fabricated a new dataset by resampling the original dataset, repeating some data-points and removing some others so that the total number of points, here microtubules, is conserved (Figure S1D, green shading). We repeated this operation for each embryo separately. We performed global fitting and obtained a new set of parameters (Figure S1D, orange shading). We repeated $M$ times (about the embryo-average total number of microtubules per embryo) this overall procedure and obtained a distribution of parameters. By assuming it was Gaussian, we extracted the corresponding average, providing an estimate $\hat{\theta}$ of the parameter value (Figure S1D, blue shading; Figure S1BC). We also got the standard deviation and easily derived the 95% confidence interval for $\theta$:

$$\hat{\theta} - se \times z_{1-\alpha/2} \leq \theta \leq \hat{\theta} - se \times z_{1-\alpha/2} \qquad (10)$$

with $z_{1-\alpha/2}$ the 1- $\alpha$ /2 quantile of the normal distribution equal to 1.96 with $\alpha$ = 0.05.

The confidence intervals obtained by these two different approaches were very similar (Figure S1BC), although they rely on different hypotheses. It reinforces the strength of our error estimates.

#### 1.3 Mapping the short- or long-lived instantaneous contacts.

The nematode zygote undergoes an asymmetric division, and its cortex is polarised through PAR proteins in a very dynamical fashion (Peglion and Goehring, 2019). In turn, the imbalance in cortical pulling forces reflects polarity (Grill *et al.*, 2003; Rodriguez-Garcia *et al.*, 2018), the asymmetric microtubule dynamics being either causative, either consecutive to this force. How the centring force coordinates to that along time is still to be found. We mapped the microtubule populations along the anteroposterior (AP) axis of the embryo. Indeed, we expected space- and time-variations of the microtubule lifetimes and proportions, within each population. We adopted the following steps: (a) to get a first estimate of the best model and its parameters, we performed the DiLiPop analysis over the whole embryo and during metaphase and anaphase (Figure S4A, step 1). (b) To ensure a large enough amount of data to discern the populations, we first estimated by simulation the sample size requirement and set the time-block width accordingly (Figure S4A, step 2). (c) We then analysed the microtubule track durations by block, hence assuming that the time and space variations of microtubule dynamics are small within a block (Figure S4A, step 3). (d) Within each region and period, using the local microtubule-dynamics parameters, each track was assigned to a population (Figure S4A, step 4). (e) Finally, we computed the spatial distribution of classified microtubule contacts for each population (Figure S4A, step 5).

Step 1: To feed the simulation of step 2, we estimated the number of microtubule populations and their characteristics by performing the DiLiPop assay in the whole embryo from early metaphase to late anaphase.

Step 2: To optimize the duration of the time-block and if necessary the extent of the regions, we estimated the minimal sample size required to resolve two populations similarly to the approach used to exclude the presence of three populations (main text, p. 7). In this instance, we set two dynamically distinct populations, with lifetimes 0.5 s and 2 s and proportions 60% and 40%. Each dataset was first composed of 25 simulated embryos to mimic the untreated condition. The recovered short-lived and long-lived lifetimes were mostly accurate in all conditions (Figure S4B2), and the double-exponential was identified as the best model whatever the sample size (Figure S4B1). Thus, 350 microtubule tracks per embryo appeared sufficient for dataset composed of 25 embryos. In contrast, we had fewer embryos in treated conditions (typically 8 to 12). We thus performed a similar assay with 8 simulated embryos per dataset. We found that at least 500 tracks per embryo were necessary to identify the number of populations correctly (Figure S5A2). While the recovered short-lived lifetime was accurate whatever the sample size, the standard error on the recovered long-lived lifetime was dependent on the population size (Figure S5A1). We concluded that 1000 tracks per embryo were necessary to get less than 10% of error on the model parameters, for dataset composed of 8 embryos.

Step 3: We separated the embryos in three biologically meaningful regions mostly corresponding to embryonic polarity as prescribed by PAR-2/3 and LET-99 protein domains. Practically, we used three regions: one spanning over 0-45% of AP axis – the anterior region – in which a low level of active force generators is present; one over 45-70% – the lateral LET-99 band – in which there are no or few active force generators; one over 70-100% – the posterior-most region – which is enriched in active force generators (Cheng *et al.*, 1995; Rose and Kemphues, 1998; Grill *et al.*, 2001; Tsou *et al.*, 2002; Colombo *et al.*, 2003; Grill *et al.*, 2003; Park and Rose, 2008; Krueger *et al.*, 2010). The duration of the time-blocks was adjusted to guarantee enough tracks per region to ensure an accurate analysis as per the above *in silico* analysis (step 2). For large sets such as the untreated condition, the time-block width was set to 60 s for the study of the three cortical regions, or 30 s when considering a single region. The achieved spatiotemporal resolution is shown in Figures 4A1-2 or 6A. In contrast, for smaller embryo sets, we used only two time-blocks, one for metaphase and the other for anaphase.

Step 4: For each microtubule, we next computed its probabilities to belong to the short-lived and long-lived populations using a mono-exponential model with corresponding parameters (Figure 1E). Each microtubule is assigned to its most probable population.

Step 5: Lastly, we generated the "*DiLiPop maps*" displaying the instantaneous contact densities for the short-lived and long-lived populations. While the classification relies on dynamics locally estimated in three regions, we divided the cortex into ten regions of equal width along the AP axis to compute the instantaneous density of contacts. The contact density was then obtained similarly to (Bouvrais *et al.*, 2018). However, in contrast to that previous work, we now get one map per population.

#### 2 Establishing the requirements to distinguish various microtubule populations.

As we investigated the two-population characteristics under various perturbations, we needed to validate that the number of collected embryos is sufficient to ensure enough statistical power and support our conclusions. In particular, we aimed to assess the accuracy of lifetimes and proportions. We used an *in silico* approach (as in main text, p. 7). We placed ourselves in conditions with few embryos, i.e. 8 simulated embryos, with 3000 tracks for each. We further considered only the conditions, where a majority of fabricated datasets led to the simulated model, here double-exponential, being the best model according to Bayesian criterion. We simulated two populations and varied one of the model parameters: the short lifetime $T_1$ (Figure S5B,C,E), the long lifetime $T_2$ (Figure S5C,D,G) or the proportion $P_1$ (Figure S5B,D,F). Overall, proportions larger than 20% and lifetime differences larger than 0.6 s ensure to distinguish the two populations with only 8 embryos (Figure S5B-D). Therefore, the DiLiPop assay enables to analyse population characteristics in a broad range of cases.
